# Transient fibrosis resolves via fibroblast inactivation in the regenerating zebrafish heart

**DOI:** 10.1101/254748

**Authors:** Héctor Sánchez-Iranzo, María Galardi-Castilla, Andrés Sanz-Morejón, Juan Manuel González-Rosa, Ricardo Costa, Alexander Ernst, Julio Sainz de Aja, Javier Langa, Nadia Mercader

## Abstract

In the zebrafish (*Danio rerio*), regeneration and fibrosis after cardiac injury are not mutually exclusive responses. Upon cardiac cryoinjury, collagen and other extracellular matrix (ECM) proteins accumulate at the injury site. However, in contrast to the situation in mammals, fibrosis is transient in zebrafish and its regression is concomitant with regrowth of the myocardial wall. Little is known about the cells producing this fibrotic tissue or how it resolves. Using novel genetic tools to mark *periostin b-* and *collagen 1alpha2* (*col1a2*)-expressing cells in combination with transcriptome analysis, we explored the sources of activated fibroblasts and traced their fate. We describe that during fibrosis regression, fibroblasts are not fully eliminated, but become inactivated. Unexpectedly, limiting the fibrotic response by genetic ablation of col1a2-expressing cells impaired cardiomyocyte proliferation. We conclude that ECM-producing cells are key players in the regenerative process and suggest that anti-fibrotic therapies might be less efficient than strategies targeting fibroblast inactivation.

**SCIENTIFIC STATEMENT:** After myocardial infarction in the mammalian heart, millions of cardiomyocytes are lost and replaced by fibrotic scar tissue. While fibrosis is persistent in adult mammals, there are some vertebrates, including zebrafish, with the capacity for regeneration. This process does not occur in the absence of fibrosis. Here we studied subpopulations of collagen producing cells and analyzed their fate after complete regeneration of the zebrafish myocardium. Our data show that fibroblasts persisted in the regenerated heart, but shut down the profibrotic program. While fibrosis could be considered as detrimental to the regeneration process, our study reveals a positive effect on cardiomyocyte proliferation. Accordingly, a fibrotic response can be beneficial for heart regeneration.

## INTRODUCTION

Cardiac fibrosis is the main cause of heart failure and other cardiovascular pathologies, including those associated with myocardial infarction (MI). Fibroblasts accumulate at the injury site, where they form an extracellular matrix (ECM)-rich scar that protects the heart from ventricular wall rupture. Fibroblasts can have pleiotropic effects after MI, producing ECM proteins and influencing the electrical coupling and contractility of cardiomyocytes.

Fibroblasts are defined as mesenchymal cells that express ECM proteins such as collagens (1). Expression of collagens, however, cannot be used as the sole criterion to define a fibroblast, since these proteins are expressed by several other cell types (1). Indeed, the search for the origin of cardiac fibroblasts during pathological fibrosis has been hindered by the lack of specific fibroblast markers (2). So far, marker genes that have been shown to be more specific for profibrotic fibroblasts include Collagen 1 alpha1 (*Col1a1*) (3) and *Periostin (Postn*) (4, 5). Lineage tracing study using the Cre/lox system showed that epicardial *Tcf21-*derived cells give rise to *Postn*-expressing activated fibroblasts in response to MI in the adult mouse (6). Moreover, ablation of *Postn^+^* cells decreased post-MI survival, confirming that fibrosis is an efficient mechanism to repair the myocardium after acute cardiomyocyte loss (6).

Unlike mammals, zebrafish hearts can fully regenerate cardiac tissue upon injury (7, 8). After controlled cryoinjury of a quarter of the zebrafish cardiac ventricle, myocardial regeneration is preceded by the accumulation of ECM at the injury site (9), indicating that a fibrotic response is not incompatible *per se* with regeneration. The origin and fate of fibroblasts in the context of heart regeneration is not well known. Possible sources of cardiac fibroblasts in the injured zebrafish heart include the epicardium and epicardial-derived cells (EPDCs) (10, 11). However, they might not be the only contributors to a fibrotic response, since collagen 1 has also been detected at the endocardial border of the injury area (IA) (12).

Mechanistically, adult cardiomyocytes re-enter the cell cycle to proliferate and give rise to *de novo* cardiomyocytes, which then repopulate the injured myocardium (8). *In vitro* co-culture experiments using murine cells indicate that fibroblasts can have a beneficial impact on cardiomyocytes by promoting their proliferation (13), but their effect on regeneration has not been reported so far. Furthermore, inhibition of the Tgf-beta signaling during zebrafish heart regeneration impairs cardiomyocyte proliferation, suggesting that fibrosis could play a role in heart regeneration (14). Since Tgf-beta also directly activates pSmad3 phosphorylation in cardiomyocytes (14), the effect of fibrosis on cardiomyocyte proliferation and heart regeneration remains an open question. Here, we examined the origin of cardiac fibroblasts in the zebrafish and characterized activated fibroblast populations in the injured heart. We also analyzed the mechanisms of fibroblast clearance during zebrafish heart regeneration. Using genetic ablation of collagen-producing cells, we show that rather than being detrimental for heart regeneration, a fibrotic response is in fact necessary for cardiomyocyte proliferation.

## RESULTS

### Endogenous fibroblasts contribute to ECM deposition in the cryoinjured zebrafish heart

A population of collagen-producing cells located between the cortical and trabecular myocardium in the zebrafish heart was identified by transmission electron microscopy (15). This region was further shown to harbor a cell population labelled in the *-6.8kbwt1a:GFP* line (hereafter termed wt1a:GFP) (16), and positive for collagen XII expression (17). To unequivocally identify them as cardiac fibroblasts, we compared the gene expression profile of *wt1a*:GFP^+^ cells with the remainder of cardiac cells using whole transcriptome analysis (Fig. 1 *A* and *B*). Pathway analysis revealed an association of *wt1a*:GFP^+^ cells with fibrosis (Fig. 1 A). Moreover, *wt1a*:GFP^+^ cells expressed 8 out of 9 genes described as the most specific mammalian cardiac fibroblast marker gene (18) (Fig. 1 *B*). We next studied whether *wt1a*:GFP^+^ cells upregulate a fibrotic gene program in response to injury. Expression of *postnb* was undetectable by mRNA *in situ* hybridization (ISH) on sections of uninjured wt1a:GFP hearts (Fig. 1 *C* and *D*; n = 3/3), but upon cryoinjury *wt1a*:GFP^+^ cells expressing *postnb* were detected surrounding the injury area (IA) (Fig. 1 *E* and *F*; n = 4/4; 66 ± 7% of *wt1a*:GFP^+^ cells expressed high levels of *postnb* mRNA of a total of 660 cells counted in 4 hearts). We next crossed *wt1a:GFP* into *col1a2:mCherry-NTR*, a line allowing to mark collagen producing cells (Fig. S1, n = 4/4). In uninjured hearts, 57 ± 8% of *wt1a*:GFP^+^ cells expressed *col1a2*:mCherry-NTR (360 cells analyzed from 5 hearts, Fig. 1 *G–K* and Q; collocalization seen in n = 5/5). At 7 days postinjured (dpi), a more intense mCherry expression was observed in 48 ± 5% of *wt1a*:GFP^+^ cells (6675 *wt1a*:GFP^+^ cells analyzed from 4 hearts, Fig. 1 *L–P* and R–S; n = 4/4). The mean fluorescence level in *wt1a*:GFP^+^ cells changed from 190 ± 50 to 1210 ± 180 A. U. (n = 4). In addition, 60 ± 4% of all *col1a2*:mCherry+ cells were *wt1a*:GFP^+^ (Fig. 1 T; n = 4/4). In sum, a large proportion of *wt1a*:GFP^+^ cells express high levels of *postnb* and *col1a2* upon injury. However, not all *col1a2+* cells in the cryoinjured heart are wt1a:GFP^+^, suggesting that other cell types also contribute to fibrosis. To unambiguously demonstrate that intracardiac *wt1a+* cells contribute to fibrosis, a *wt1a:CreER^T2^* line was generated and crossed into *ubb:loxP-GFP-loxP-STOP-mCherry (ubb:Switch)*. Recombination was induced by 4-Hydroxytamoxifen (4-OHT) administration at 4 and 3 days prior to injury (Fig. S2 *F*). In uninjured hearts, *wt1a*-derived cells were observed between the trabecular and cortical myocardium, colocalizing with low levels of col1a1 immunofluorescence staining (Fig. S2 *G–J*; n = 4/4). By contrast, in cryoinjured hearts, *wt1a*-derived cells were surrounded by a strong signal of col1a1 immunostaining (Fig. S2 *K–N;* n = 4/4). Furthermore, recombination in the double transgenic line *wt1a:CreER^T2^*; *col1a2:loxP-tagBFP-loxP-mCherry-NTR* allowed mCherry labeling of cells close to the IA (Fig. S2 *O–S*; n = 3/3). In the mouse and the chicken, intracardiac fibroblasts have been described to derive mainly from the embryonic epicardium (19). We therefore lineage traced embryonic *wt1a-*derived cells in the zebrafish (Fig. S2 *A*). In the embryo, *wt1a*:GFP marks a population of proepicardial cells (16). Recombination in two-day-old embryos yielded labeled epicardial cells on the surface of the myocardium at 5 days postfertilization (dpf) (Fig. S2 *B–E*; n = 8/8). In the adult, while some embryonic *wt1a*-derived cells remained within the epicardial layer, they could also be observed in deeper cell layers within the myocardium (Fig. S2 *F–I*; n = 3/3). In line with single cell analysis of *tcf21*-derived cells in zebrafish, this suggests that intracardiac fibroblasts in the zebrafish also derived from the embryonic epicardium (20). To confirm that EPDCs contribute to cardiac fibrosis, we crossed *tcf21:CreER^T2^* into *col1a2:loxP-tagBFP-loxP-mCherry-NTR*, to label the *col1a2*^+^ cells that expressed *tcf21* at the time of 4-OHT administration (Fig. S2 *J*). The presence of mCherry in injured hearts confirmed that *tcf21*-derived cells contribute to collagen deposition (Fig. S2 *K–N*; n = 5/5). Overall, these results show that EPDC-derived resident cardiac fibroblasts secrete collagen in response to injury in the zebrafish heart.

**Fig 1.**
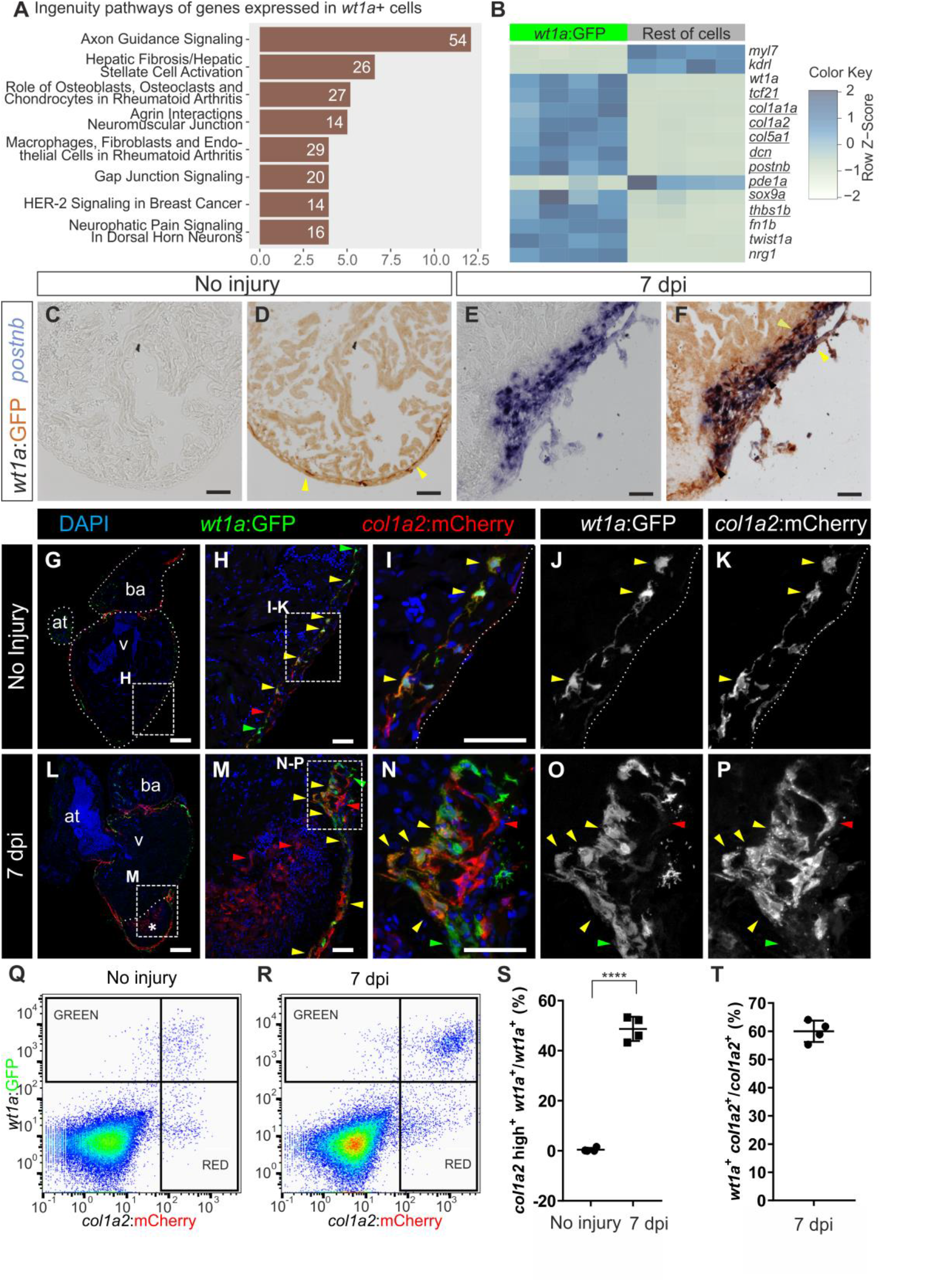
Intracardiac fibroblasts contribute to transient fibrosis during zebrafish heart regeneration. (*A* and *B*) Transcriptome analysis of *wt1a*:GFP^+^ cells isolated from adult zebrafish hearts: (*A*) Ingenuity pathways enriched in *wt1a*:GFP^+^ ventricular cells compared with all GFP^-^ cells. Number of differentially expressed genes are shown; x axis shows –log10(B.H.-p-values. (*B*) Heat map indicating upregulation of fibrotic marker genes (underlined) in the *wt1a*:GFP^+^ cell fraction. *(C–F) postnb* mRNA *in situ* hybridization followed by anti-GFP immunohistochemistry on sections of *wt1a*:GFP ventricles without injury (*C* and *D*) and at 7 days postinjury (dpi) (*E* and *F*). Arrowheads, *wt1a*:GFP^+^ cells. (*G–P*) Immunofluorescence staining on sections of *wt1a:GFP; col1a2:mCherry-NTR* double transgenic hearts without injury (*G–K*) or at 7 dpi (*L–P*). Red arrowheads, *col1a2*:mCherry^+^ cells; green arrowheads, *wt1a*:GFP^+^ cells; yellow arrowheads, double positive cells. (*Q* and *R*) FACs-sorted cells from *wt1a:GFP;col1a2:mCherry* ventricular apex without injury (*Q*) or at 7 dpi (*R*). Representative examples from a total of 4 hearts per condition analyzed. (*S*) Percentage of *wt1a*:GFP^+^ cells that express more than 1000 A.U. of mCherry, threshold corresponding to the maximum value of mCherry detected in non-activated fibroblasts. (*T*) Percentage of *col1a2*:mCherry^+^;*wt1a*:GFP^+^ cells according to the thresholds shown in panel *R*. Graphs show individual measurements and means ± s.d. **** *P* -*--< 0.0001 by two-tailed t-test. Scale bars, 25 μm (*H,I,M,N*), 50 μm (*C–F*), 100 μm (*G* and *L*).

### Kidney marrow-derived cells do not contribute to cardiac fibrosis in the zebrafish

We next assessed whether cells derived from the main hematopoietic organ in the zebrafish, the kidney marrow, contribute to cardiac fibrosis in the zebrafish. Kidney marrow-derived cells (KMDCs) from *ubb:GFP; col1a2:mCherry-NTR* fish were transplanted into irradiated wildtypes to reconstitute the hematopoietic system with fluorescently labeled cells (Fig. S3 *A*). After heart cryoinjury and fixing at 7 dpi GFP^+^ KMDCs colonized the injured heart (n = 5/5). No mCherry expression could be detected in the hearts at 7 dpi (Fig. S3 *B–E*; n = 5/5), a stage in which *col1a2*:mCherry expression was readily observed in a control group of donors (Fig. S3 *F–I*; n=4/4). This finding suggests that KMDCs do not contribute to ECM production during cardiac fibrosis in the zebrafish.

### Endocardial cells have a limited contribution to ECM deposition in the cryoinjured zebrafish heart

To further characterize the contribution of the endocardium to fibrosis we analyzed the gene expression signature of endocardial cells in response to injury. We utilized the *kdrl:mCherry* transgenic line, which predominantly labels endocardial cells and relatively few coronary arteries (21) and performed RNA-Seq analysis of fluorescence activated cell sorter (FACS)-sorted mCherry^+^ cells from the apical region of control hearts and the injured ventricular apex of hearts at 7 dpi (Fig. 2 A–B). The most upregulated genes in *kdrl*^+^ cells were related to fibrosis. To validate the RNA-Seq results, we crossed the endothelial and endocardial reporter line *fli1a:GFP* with *col1a2:mCherry-NTR* fish. At 7 dpi, expression of GFP and mCherry colocalized at the IA, but not in the border zone (Fig. 2 *C–F*; n = 5/7). Cell sorting revealed that upon injury, a small proportion of *fli1a*:GFP^+^ cells expressed high levels of mCherry (Fig. 2 *G–I*): 19 ± 14% of *fli1a*:GFP^+^ cells were *col1a2*:mCherry-NTR+ (Fig. 2 *J*). Next, *fli1a*^+^ were traced with a newly generated *fli1a:CreER^T2^* line crossed into *ubb:Switch*, (Fig. S4 *A*). In uninjured hearts, mCherry^+^ cells lined the endocardial border, revealing efficient recombination (Fig. S4 *B–E*; n = 5/5). At 7dpi, *fli1a*-derived cells were found in close proximity to col1a1 staining at the IA (Fig. S4 *F–I*). We also crossed *fli1a:CreER^T2^* into the *col1a2:loxP-tagBFP-loxP-mCherry-NTR* line (Fig. 2 *K*). At 7 dpi, mCherry^+^ cells were present at the IA, but not the periphery, revealing that *fli1a*-derived cells produce collagen locally at the IA (Fig. 2 *L–O*; n = 7/12). Although endocardial cells close to the IA become more rounded, they did not appear to adopt a mesenchymal phenotype and remained connected, forming a layer around the luminal border of the site of injury. In agreement, while epithelial-mesenchymal transition (EMT) markers were upregulated in *wt1a*:GFP^+^ cells upon injury, *kdrl*:mCherry^+^ cells revealed low levels of *twist1a, prrx1a* and *snail2* expression(Fig. S4 *J–L*).

**Fig. 2.**
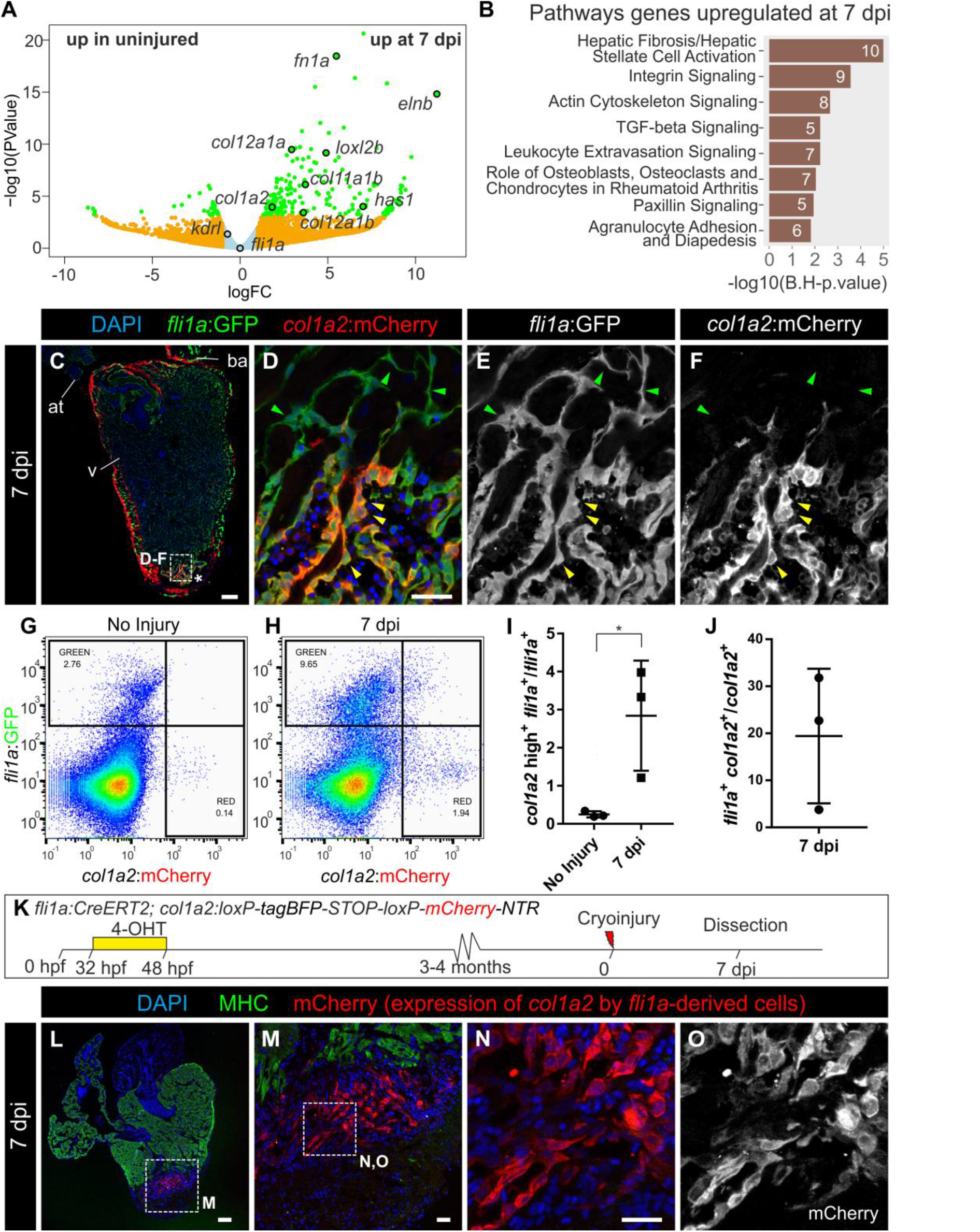
Endocardium-derived cells contribute to transient fibrosis during zebrafish heart regeneration. (*A*) The transcriptome of *kdrl*:mCherry^+^ cells FACs-sorted from the ventricular apex of hearts with no injury or at 7 dpi was performed. Volcano plot. Light blue, false discovery rate (FDR) > 0.05, abs(log fold change (LFC)) < 1; orange, FDR > 0.05, abs(LFC) > 1; red, FDR < 0.05, abs(LFC) < 1; green, FDR < 0.05, abs(LFC) > 1. (B) Ingenuity pathway analysis including the number of differentially expressed genes.(*C–F*) Immunofluorescence with anti-GFP and anti-mCherry on a heart section from an adult *fli1a:GFP;col1a2:mCherry-NTR* zebrafish. Yellow arrowheads: double-positive cells at the injury area (IA; asterisk). Green arrowheads: *fli1a:GFP*^+^ cells negative for mCherry. (*G–H*) FACs-sorted cells from *fli1a:GFP;col1a2:mCherry* hearts without injury or at 7 dpi. Shown are 2 representative examples from a total of 4 hearts per condition analyzed. (*I*) Percentage of *fli1a:GFP*^+^ cells that express more than 100 A.U. of mCherry, threshold corresponding to the maximum value of mCherry detected in endocardial cells from uninjured hearts. (*J*) Percentage of *col1a2:mCherry*^+^; *fli1a:GFP*^+^ cells according to the thresholds in panel (*H*). Graphs show individual measurements form hearts as well as mean ± s.d. values. \**P* = 0.0363 by two-tailed t-test. (*K*) Experimental scheme for visualizing collagen-producing endocardial cells. (*L–N*) Immunostaining of a heart section close to the IA: (*M*) zoomed view of (*L*). (*N* and *O*) zoomed views of (*M*). mCherry marks *fli1a*-derived cells expressing *col1a2*, MHC marks the myocardium and nuclei are DAPI counterstained. at, atrium; v, ventricle. Scale bars, 25 μm (*D, M, N*), 100 μm (*C, L*).

### *Postnb* expression reports activated fibroblasts in the injured zebrafish heart

To characterize *postnb* expressing cells that appear in response to injury, we generated a *postnb*:citrine reporter line. In adult uninjured hearts, *postnb*:citrine expression was detected in the bulbus arteriosus, valves, some perivascular cells associated with large coronary arteries in the basal ventricle, and in isolated epicardial cells (Fig. 3 *A–C*; n = 3/3), but not in the ventricular apex (Fig. 3 *B*). At 7 dpi, *postnb*:citrine^+^ cells were surrounding the IA in the cryoinjured apex and exhibited a mesenchymal morphology (Fig. 3 *D* and *E*; n = 3/3 and Fig. 3 *F* and *G*; n = 7/7). *col1a2*^+^ and *postnb*^+^ cells did not colocalize completely (Fig. S5 *A* and *B*; n = 3/3 hearts, and *C–G*; n = 6/6 hearts). Moreover, we did not observe cells co-expressing postnb:citrine and *fli1a*:DsRed or *kdrl*:mCherry (Fig. 3 *F* and *G* n = 4/4 and Fig. 3 *I*; n = 5/5), further confirming that at least some of the collagen producing cells have an endocardial origin. The *postnb*^+^ population expanded during the first week postinjury, as revealed by proliferator cell nuclear antigen (PCNA) staining in *postnb*^+^cells sampled at 7 and 14 dpi (Fig. 3 *H*). After 3 weeks, the number of *postnb*/PCNA double-positive cells significantly decreased. To further characterize this population, we analyzed the transcriptome of *postnb*:citrine^+^ cells sorted from the injured ventricle, identifying pathways related to fibrosis and axon guidance (Fig. 3 J). Compared with the remainder of cells at the IA, *postnb*:citrine^+^ cells were highly enriched for genes encoding secreted proteins. In total, 128 out of 917 differentially expressed genes upregulated in the postnb:citrine+ population encoded secreted molecules, while only 12 out of 2206 enriched in the negative population belonged to this category (*P* < 0.00001 by chi-square test). *postnb:citrine*^+^ cells expressed several ECM proteins in addition to 12 matrix metalloproteases (Fig. 3 *K* and Fig. S5 *H-I*). Interestingly, *postnb:citrine*^+^ cells also expressed several signaling molecules (Fig. 3 *K* and Fig. S5 *J*). While some of these molecules have been described to influence heart development or regeneration (22–26), several others have not been studied in the context of heart regeneration to date, including *wnt5a, wnt16, rspo1* and *hhip* (Fig 3 *K*).

**Fig. 3.**
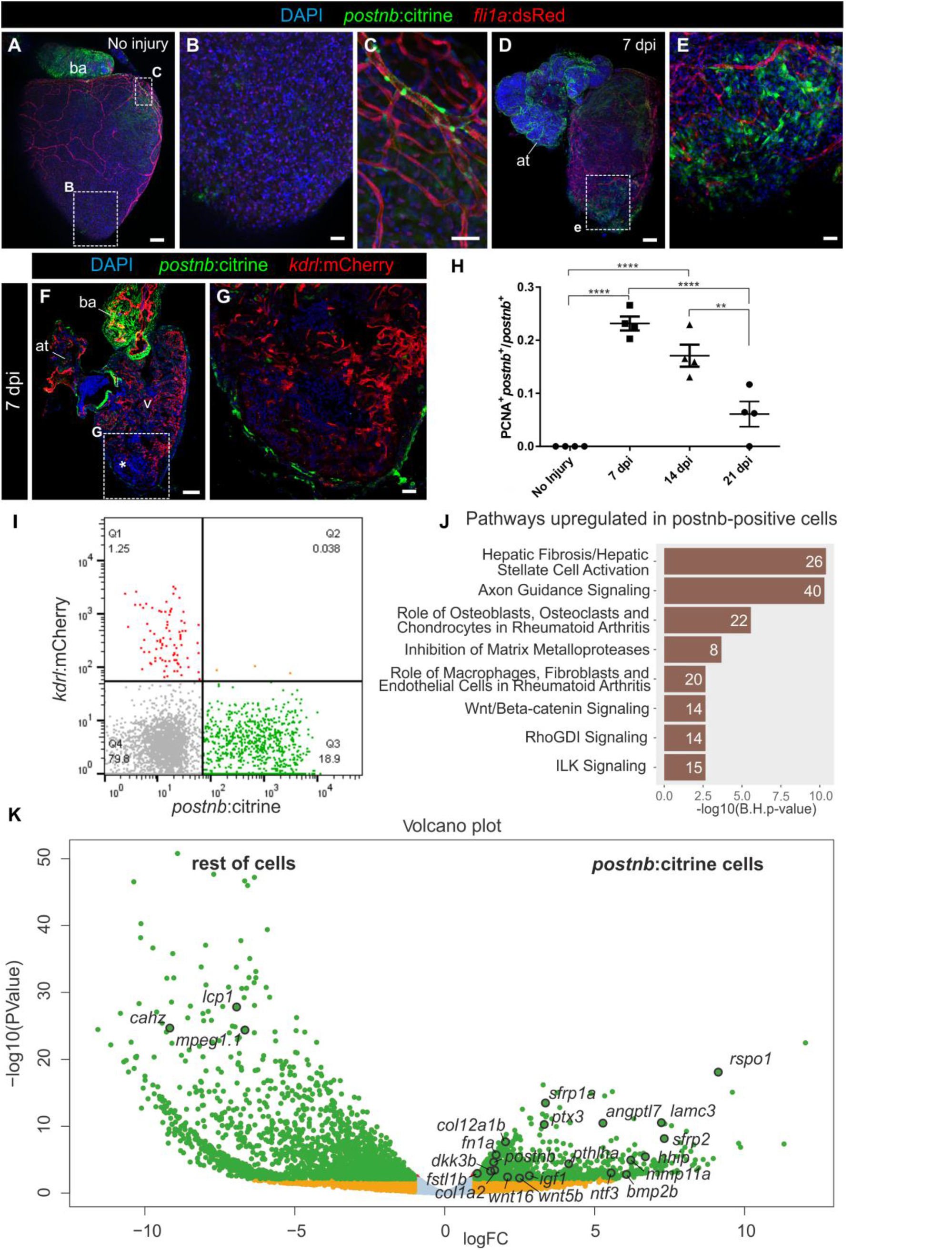
*periostin b* expression marks an activated cardiac fibroblast population upon ventricular cryoinjury. (*A–E*) Whole-heart immunofluorescence in the *postnb:citrine; fli1a:dsRedEx* double transgenic line. (*A*) Whole-heart and (*B* and *C*) zoomed views of the ventricular apex of an uninjured heart and a heart at 7 dpi (*D* and *E*). Perivascular cells can be observed in (*C*). *postnb*:citrine, green; *fli1a*:dsRedEx, red. (*F* and *G*) Immunofluorescence staining on a sagittal heart section of a *postnb*:citrine; *kdrl*:mCherry zebrafish (*F*). (*G*) Zoomed view of the injured ventricular apex (asterisk); *postnb*:citrine, green; *kdrl*:mCherry, red. (*H*) Quantification of proliferating *postnb*^+^ cells postinjury (mean ± s.d; ^****^*P* < 0.0001; ^**^*P* < 0.01 by one-way ANOVA followed by Tukey's multiple comparisons test). (*I*) FACs-sorted cells from *kdrl*:mCherry;*postnb*:citrine hearts. No double-positive cells were detected. (*J* and *K*) Transcriptome analysis of *postnb*:citrine^+^ cells isolated from the ventricular apex. (*J*) Ingenuity pathway analysis, x axis shows –log10(B.H.-p-values). (*K*) Volcano plot. Light blue, false discovery rate (FDR) > 0.05, abs(log fold change (LFC)) < 1; orange, FDR > 0.05, abs(LFC) > 1; red, FDR < 0.05, abs(LFC) < 1; green, FDR < 0.05, abs(LFC) > 1. at, atrium; ba, bulbus arteriosus; v, ventricle. Scale bars, 25 μm (*B, C, E, G*), 100 μm (*A, D, F*).

Overall, these results reveal that the *postnb:citrine* zebrafish line marks activated fibroblasts, which express gene encoding ECM proteins as well as proteins responsible for ECM degradation. Furthermore, they express secreted signaling genes that could influence heart regeneration.

### Fibroblast inactivation leads to fibrosis regression during cardiac regeneration

Whereas cardiac fibrosis is irreversible in adult mammals, degradation of fibrotic tissue occurs in the zebrafish, and the expression of fibrosis-related genes drops to near baseline levels at 90 dpi (Fig. S6 *A* and *B* and (27)). Elimination of activated fibroblasts has been suggested to progress via a mechanism of programmed cell death (1). However, Terminal deoxynucleotidyl transferase dUTP nick end labeling (TUNEL) staining on *postnb*:citrine heart sections at different postinjury stages revealed a low number of apoptotic *postnb*^+^ cells at 7 and 14 dpi and none at 21 dpi (Fig. S6 *C–G*). Given the low number of apoptotic fibroblasts identified at the 3 stages analyzed, fibroblasts might remain in the heart even after complete regeneration. To test this, we generated a *postnb:CreER^T2^* line and crossed it with the *ubb:Switch* line (Fig. 4 *A*). Two pulses of 4-OHT at 3 and 4 dpi allowed permanent mCherry labeling of *postnb*^+^ cells. *postnb*-derived cells were detected at the IA at 7 dpi (Fig. 4 *B–E*; n = 4/4). Surprisingly, at the ventricle apex, *postnb*-derived cells could still be detected within the regenerated myocardium at 90 dpi, albeit in lower numbers (Fig. 4 *F–J*; n = 5/5). To assess whether fibroblasts revert to a quiescent phenotype during regeneration, we performed RNA-Seq analysis of *postnb*-derived cells at 7 and 60 dpi (Fig. 4 *K–N*). Pathway analysis revealed differences in the expression profiles of *postnb*-derived cells between these stages (Fig. 4 *K* and *L*). Although overall the level of ECM gene expression decreased from 7 to 60 dpi, there were also collagen-encoding genes that were upregulated at 60 dpi, such as *col7a1l* and *col8a2* (Fig. 4 *M*–N and Fig. S7 *A–I*). These genes were not expressed by *wt1a*^+^ resident fibroblasts in the uninjured heart, indicating that the inactivation of fibroblasts does not fully revert the expression profile to a homeostatic baseline. Overall, the gene signature of postnb-derived cells at 60 dpi resembled more the signature of *wt1a*:GFP cells than that of postnb-derived cells at 7 dpi (Fig. 5 *O* and Fig. S7 *J–L*). Indeed, *postnb*^-^ *wt1a*:GFP^+^ cells were found at 130 dpi in the regenerated myocardium (Fig. S8).

**Fig. 4.**
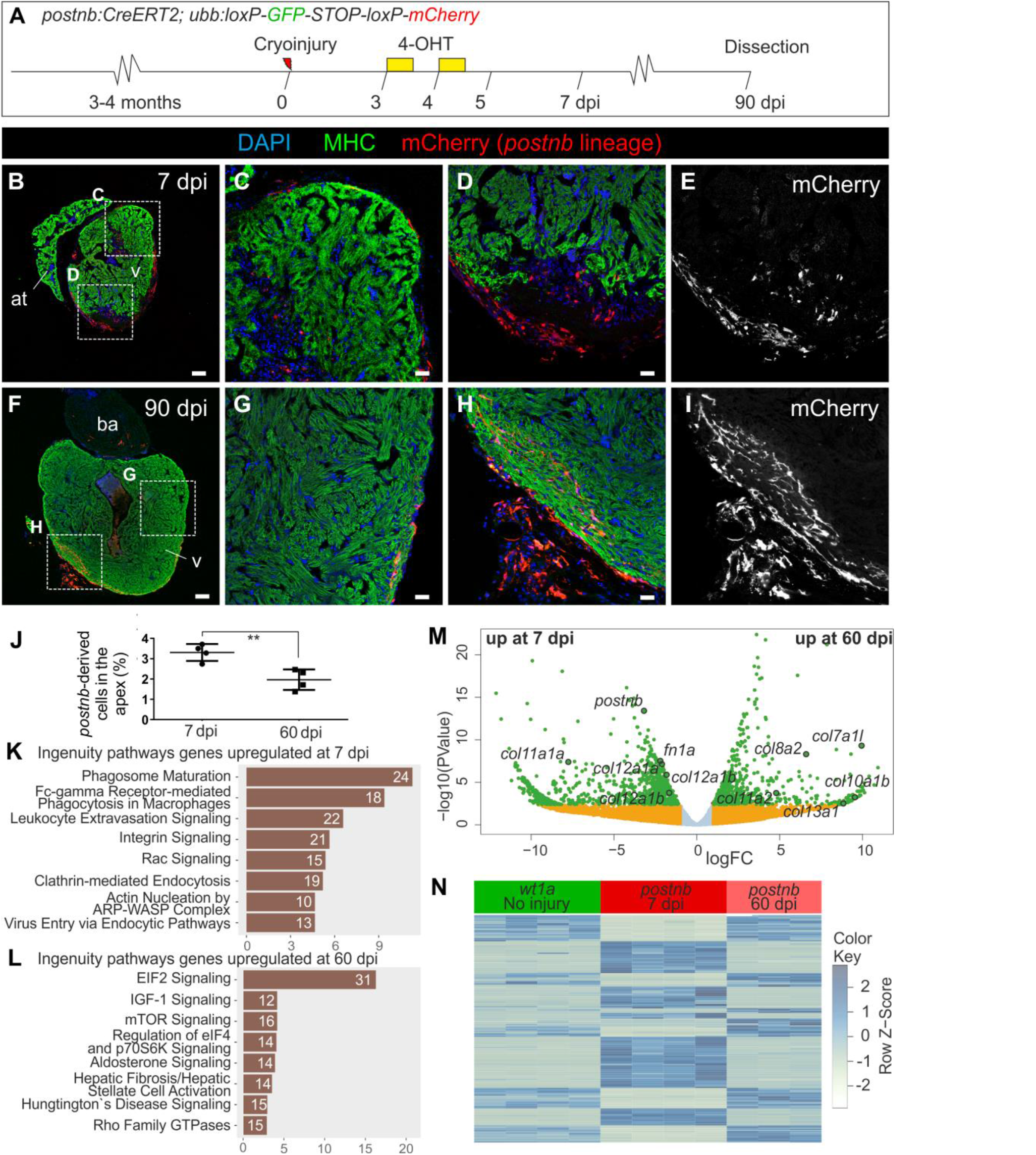
Persistence of *postnb*-derived cells in the regenerated myocardium. (*A*) 4-Hydroxytamoxifen (4-OHT) was added to *postnb*:CreER^T2^;*ubb*:Switch fish at 3 and 4 dpi, and hearts were dissected at different dpi. (*B–I*) Immunofluorescence staining on heart sections at 7 (*B–E*) or 90 dpi (*F–I*). (*J*) Percentage of *postnb*-derived cells at the injury area. Symbols show individual measurements and boxes and whiskers show mean ± s.d.; \*\**P* = 0.0064 by two-tailed unpaired t-test. (*K–M*) postnb-derived mCherry^+^ cells were sorted from the ventricular apex at 7dpi and 60 dpi, and transcriptome analysis was performed on isolated mCherry^+^ cells. (*K* and *L*) Ingenuity pathway analysis. Bars represent pathways enriched in *postnb*-derived cells compared with the remainder of cells in the injury area at 7 and 60 dpi. Numbers of differentially expressed genes are indicated. x axis shows –log10(B.H.-p-values). (*N*) Volcano plot. Light blue, false discovery rate (FDR) > 0.05, abs(log fold change (LFC)) < 1; orange, FDR > 0.05, abs(LFC) > 1; red, FDR < 0.05, abs(LFC) < 1; green, FDR < 0.05, abs(LFC) > 1. (*M*) Heat map of the top 200 differentially expressed genes between 7 and 60 dpi in *postnb*-traced cells and their expression in *wt1a*:GFP^+^ cells from uninjured hearts. at, atrium; ba, bulbus arteriosus; dpi, days post injury; MHC, myosin heavy chain; v, ventricle. Scale bars, 25 μm (*C, D, E, G, H, I*), 100 μm (*B* and *F*).

**Fig 5.**
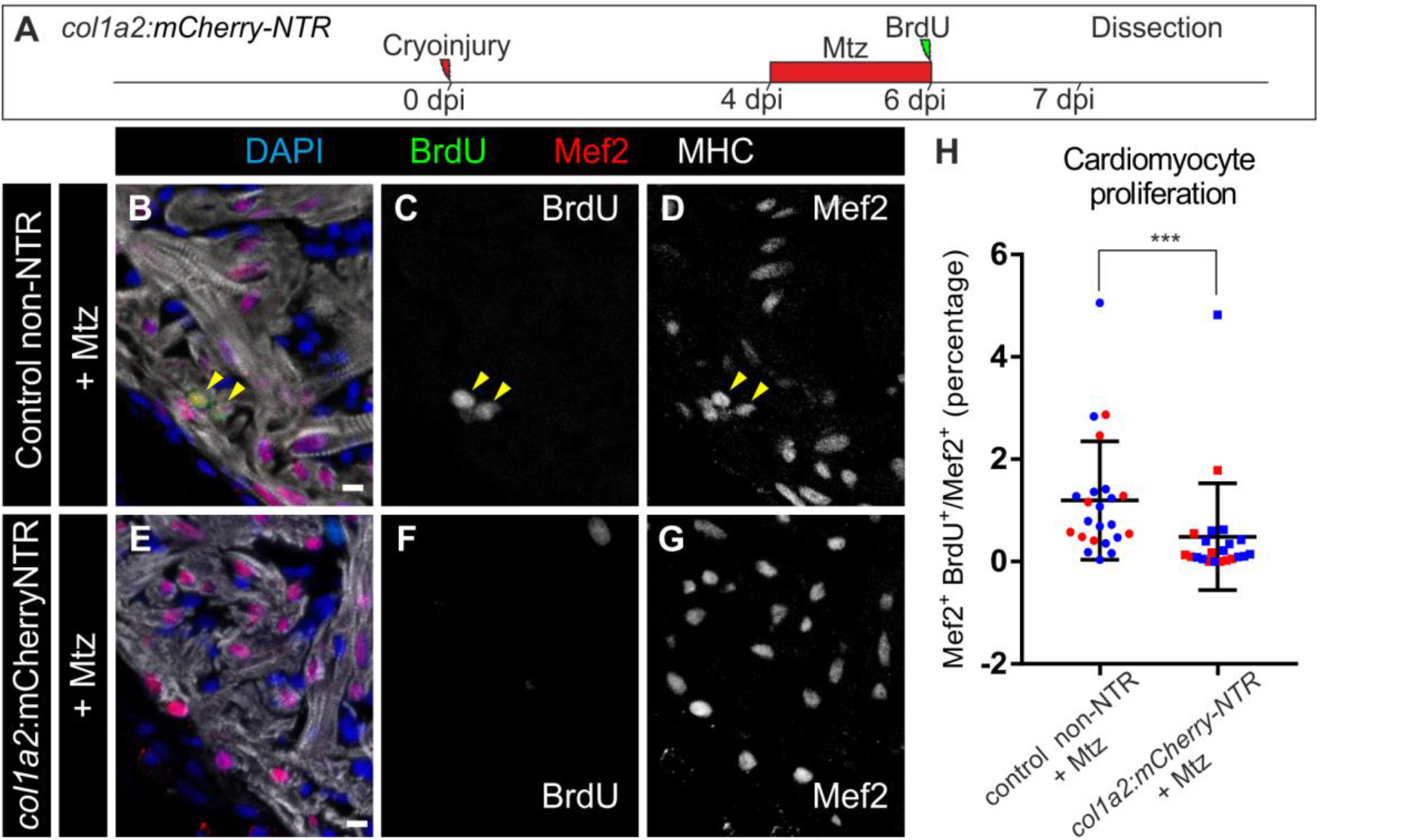
Genetic ablation of *collagen 1a2* expressing cells impairs cardiomyocyte proliferation in the cryoinjured heart. (*A*) *col1a2:mCherry-NTR* adult animals were cryoinjured and treated with Metronidazole (Mtz) from 4 to 6 dpi. BrdU injection was performed one day prior to fixation. (*B–G*) Immunofluorescence using anti-mef2 and anti-MHC to mark cardiomyocytes, and anti-BrdU in *col1a2:loxP-tagBFP-loxP-mCherry–NTR* (control) and *col1a2:mCherry-NTR* fish treated with Mtz and BrdU as described in (*A*). (*H*) Quantification of BrdU^+^ cardiomyocytes in *col1a2:mCherry-NTR* and control hearts. Shown are individual measurements and median ± interquartile range; \*\*\**P* = 0.0004 by Mann-Whitney test, n = 23 fish per condition, from 2 different experiments (blue and red dots). 3 whole-heart ventricle sections were quantified per point. NTR, Nitroreductase; MHC, myosin heavy chain. Scale bars, 10 μm (*C, E, F, I*), 100 μm (*B* and *D*).

Altogether, the data indicate that during fibrosis regression, activated fibroblasts partially return to a quiescent stage without fully resembling endogenous cardiac fibroblasts from the uninjured heart.

### Fibroblasts influence cardiomyocyte proliferation during heart regeneration

To test whether cardiac fibrosis influences heart regeneration, we ablated *col1a2*-expressing cells using a *col1a2:mCherry-NTR* line, in which *col1a2*^+^ cells express nitroreductase (NTR) and are ablated upon Metronidazole (Mtz) treatment (Fig. S9 A). Fragmented *col1a2*^+^ cell bodies and nuclei were found in Mtz-treated animals at 7 dpi, indicating efficient ablation (Fig. S9 *B* and *C*; n = 8/8). This was not detected in the untreated control group (Fig. S9 *D* and *E*; n = 3/3). No significant differences in heart regeneration were observed between the experimental and control groups at 35 dpi, suggesting that ablation of ECM-producing cells does not enhance regenerative capacity (Fig. S9 *F-M*). Nonetheless, collagen 1 could still be detected upon genetic ablation (Fig. S10). However, we found a four-fold reduction in cardiomyocyte proliferation in *col1a2:mCherry-NTR* fish compared with control siblings (Fig. 5 *A–H*).

In sum, our results not only suggest that fibrosis is compatible with regeneration, but also indicate that transient fibrosis upon cryoinjury is necessary for cardiomyocyte proliferation.

## DISCUSSION

In the mouse, cardiac repair upon MI or ventricular overload has been shown to occur mainly by ECM deposition from intracardiac fibroblasts (3, 6, 28, 29) and the epicardium (30, 31), whereas a contribution from the bone marrow is controversial (3, 28, 32-37). Endothelial cells have been suggested to contribute to cardiac fibrosis in a pressure overload model (3, 28, 36). Our data reveal that in the context of heart regeneration, mostly pre-existing fibroblasts but also endocardial cells, contribute to collagen production. Co-expression of *fli1a*^+^ or *fli1a*-derived cells and *col1a2* was not observed in all analyzed hearts, suggesting that the endocardium is not the principal contributor to fibrosis. Of note, endocardial cells at the injury border failed to undergo full EMT, and thus do not adopt a complete fibroblast-like phenotype. Cells from the epicardial border were found to produce both periostin and collagen, whereas the endocardial cells only produce collagen. These two distinct ECM environments surrounding the injured area may play an important role to guide heart regrowth.

In contrast to mammals, where ECM persists after MI, it is degraded in the zebrafish heart. We found that downregulation of fibroblast ECM production is a key step for fibrosis regression. Interestingly, clearance does not involve the complete elimination of ECM-producing cells. Indeed our data suggest that *postnb*^+^ cells might not only be actively involved in ECM production, but may also participate in its degradation during regeneration.

*tcf21* -derived cells detected in the regenerated myocardium are epicardial cells or EPDCs that were already present in the uninjured myocardium, and whether they represent inactivated fibroblasts, has not been previously explored (38). Our results suggest that this population represents *tcf21*^+^ cells that activated *postnb* in response to injury. The herein reported long-term permanence of *postnb*-derived cells in the regenerated heart might also provide an explanation for why there is only a partial recovery of ventricular wall contraction after cryoinjury (39). Our data also demonstrate that ablation of ECM-producing cells at an early stage of the injury response impairs cardiomyocyte proliferation. Thus, fibrosis has a beneficial effect on heart regeneration. Interestingly *col1a2*^+^ cells affect cardiomyocyte proliferation through mechanisms not directly linked to collagen deposition. *postnb*-derived cells express several secreted molecules that promote cardiomyocyte proliferation in the zebrafish. Furthermore, heart regeneration requires heart re-innervation (40, 41), and our gene expression analysis suggests that fibroblasts might also contribute to axon pathfinding. While cardiomyocyte proliferation was impaired, cardiac regeneration recovered after the ablation of collagen-producing cells. This may suggest compensatory mechanisms, or indicate that a recovery occurring after ablation can overcome the delayed regeneration.

In the adult mouse, ablation of Postn-derived cells impairs the recovery of cardiac pumping efficiency upon MI, revealing an important role for fibroblasts during cardiac repair (6). The lack of fibroblasts was proposed as an explanation for the regenerative capacity of the zebrafish heart (42). However, here we confirm the presence of cardiac fibroblasts in the zebrafish, and reveal that they not only contribute to the fibrotic response after injury, but are also necessary for cardiomyocyte proliferation during regeneration. This new information on how fibrosis influences myocardial regeneration and how it is eliminated in a species with endogenous regenerative potential could have important implications for regenerative medicine strategies.

## MATERIALS AND METHODS

Experiments were approved by the Community of Madrid “Dirección General de Medio Ambiente” in Spain and the “Amt für Landwirtschaft und Natur” from the Canton of Bern, Switzerland. Animals were housed and experiments performed in accordance with Spanish and Swiss bioethical regulations for the use of laboratory animals. Animals used for experiments were randomly selected from siblings of the same genotype. Detailed information on the lines used and generated can be found in the Supplementary Material and Methods sections.

ISH on paraffin sections and on whole mount larvae was performed as described (10, 43). A detailed protocol and list of riboprobes and antibodies used for immunofluorescence staining can be found in the supplementary information text file.

Apoptosis was detected by TUNEL staining using the *in situ* cell death detection kit from Roche (Mannheim, Germany).

A Leica TCS SP-5 confocal microscope was used for immunofluorescence imaging on sections and whole mount hearts, whereas a Nikon 90i was used to image non-fluorescent sections. Images were quantified manually in ImageJ (NIH) using the Cell Counter plugin.

RNA-Seq data have been deposited in GEO with the accession numbers GSE101204, GSE101200 and GSE101199. A detailed protocol can be found in Supplementary Material and Methods sections.

## ACKNOWLEDGEMENTS

We are grateful to the Animal Facility, Histology, Microscopy, Cellomics, Bioinformatics and Genomics Units from CNIC, and the Microscopy Imaging Center of the University of Bern. We thank C. Helker for sharing reagents, and O. Kanisikak for discussion. H.S and N.M. are funded by the Spanish Ministry of Economy and Competitiveness (MINECO) by grants FPU12/03007 and BFU2014–56970–P, Plan Estatal 2013-2016, Programa Estatal de I+D+i: Proyectos I+D+i 2016, and Fondo Europeo de Desarrollo Regional. N.M. is supported by the Swiss National Science Foundation grant 31003A_159721 and the ERC starting grant 337703–zebra–Heart. A.E. is funded through ANR-SNF Project 320030E-164245 to N.M. The CNIC is supported by MINECO and the ProCNIC Foundation, and is a Severo Ochoa Center of Excellence (MINECO award SEV-2015-0505).

## AUTHOR CONTRIBUTION

H.S. performed the experiments and contributed to the experimental design. M. G.-C., A. S.-M., J.M.-G.R., R.C., A.E., J.G. and J.L. contributed to experiments. N.M. designed experiments, wrote the manuscript and secured funding. All authors contributed to writing the manuscript.

## SUPPLEMENTARY INFORMATION

### SUPPLEMENTARY FIGURES

**Figure S1.**
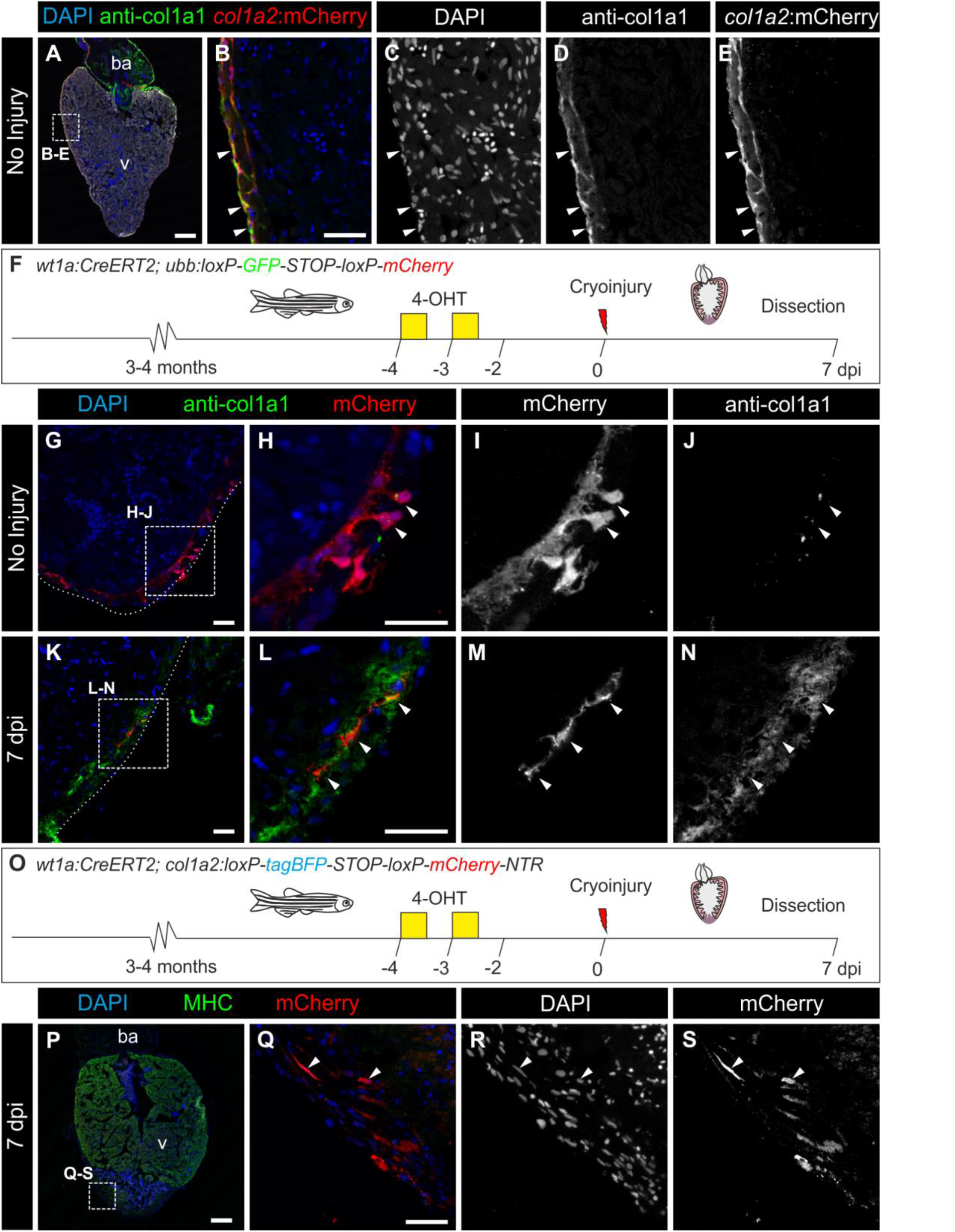
The *col1a2:mCherry-NTR* transgenic line allows the labeling of collagen producing cells and wt1a-derived cells express *col1a2* after cryoinjury. (*A*) Immunofluorescence on heart sections with anti-col1a1 (green) and mCherry (red). Nuclei are counterstained with DAPI. (*B* and *E*) Merged and individual channels of the boxed area in (*A*). Arrowheads mark mCherry^+^ cells that are surrounded by col1a1. (*F–N*) Lineage tracing of *wt1a*^+^ cells. (*F*) 4-Hydroxytamoxifen (4-OHT) was added to adult *wt1a:CreER^T2^; ubb:Switch* uninjured fish 10 and 9 days before dissection. (*G–N*) Immunofluorescence staining with anti-col1a1 (green) and mCherry (red) on heart sections of uninjured hearts (*G–J*) or 7 days post injury (dpi) hearts (*K–N*). Nuclei are DAPI counterstained (blue). Arrowheads mark wt1a-derived cells. (*O*) Experimental scheme for tracing the fate of *wt1a*-derived cells expressing *col1a2*. The *wt1a:CreER^T2^* line was crossed into the *col1a2:loxP-tagBFP-STOP-loxP-mCherry-NTR* line, in which mCherry-NTR is not expressed. Upon 4-OHT administration, recombination of loxP sites leads to activation of mCherry expression under the control of a *col1a2* promoter. Hearts from animals at 7 days postinjury (dpi) were dissected and sectioned. (*P–S*) Immunofluorescence on heart sections with anti-Myosin Heavy Chain (MHC, green), and mCherry (red). Nuclei are counterstained with DAPI. (*Q-S*) Merged and individual channels of the boxed area in P. Arrowheads mark mCherry^+^ cells, which express *col1a2*. ba, bulbus arteriosus; v, ventricle. Scale bars, 25 m (*B, G, H, K, L, Q*), 100 m (*A, P*).

**Figure S2.**
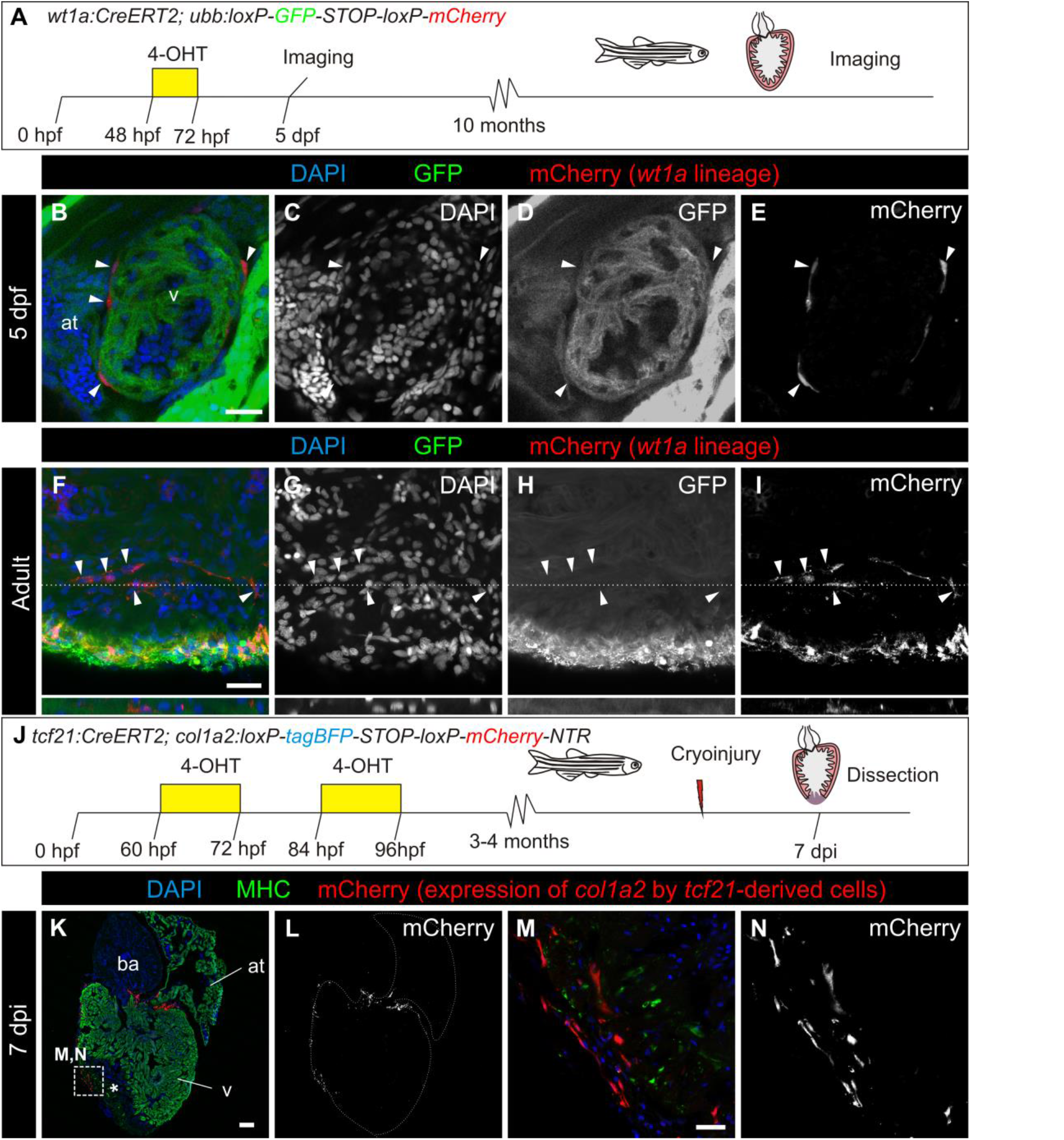
Resident fibroblasts are derived from the epicardium, and cells derived from epicardium and resident fibroblasts express *col1a2*:mCherry after injury. (*A*) Experimental scheme for tracing the fate of *wt1a*-derived cells. 4-Hydroxytamoxifen (4-OHT) was administered from 48 to 72 hours post fertilization (hpf). (*B–E*) Immunofluorescence of 5 days post fertilization (dpf) old embryos with anti-GFP (green) and mCherry (red). Nuclei are counterstained with DAPI. (*C–E*) are single channels of merged image shown in (*B*). (*F–I*) Immunofluorescence of whole mount adult hearts with anti-GFP (green) and mCherry (red). Nuclei are counterstained with DAPI. Orthogonal views of the plane highlighted with dotted lines are shown below. Shown are single (*G–I*) and merged (*F*) channels. (*J*) Experimental scheme for tracing the fate of *tcf21*-derived cells expressing *col1a2*. The *tcf21:CreER^T2^* line was crossed into the *col1a2:loxP-tag BFP-STOP-loxP-mCherry-NTR* line. Upon 4-OHT administration, recombination of loxP sites leads to activation of mCherry expression under the control of a *col1a2* promoter. Hearts from animals at 7 dpi were dissected and sectioned. (*K–N*) Immunofluorescence on the heart sections with anti-Myosin Heavy Chain (MHC, green), and mCherry (red). Nuclei are counterstained with DAPI. (*M* and *N*) Merged and individual channels of the boxed area in (K). Arrowheads mark mCherry^+^ cells, which express *col1a2*. at, atrium; ba, bulbus arteriosus; v, ventricle. Scale bars, 25 μm (*B, F, M*), 100 μm (*K*).

**Figure S3.**
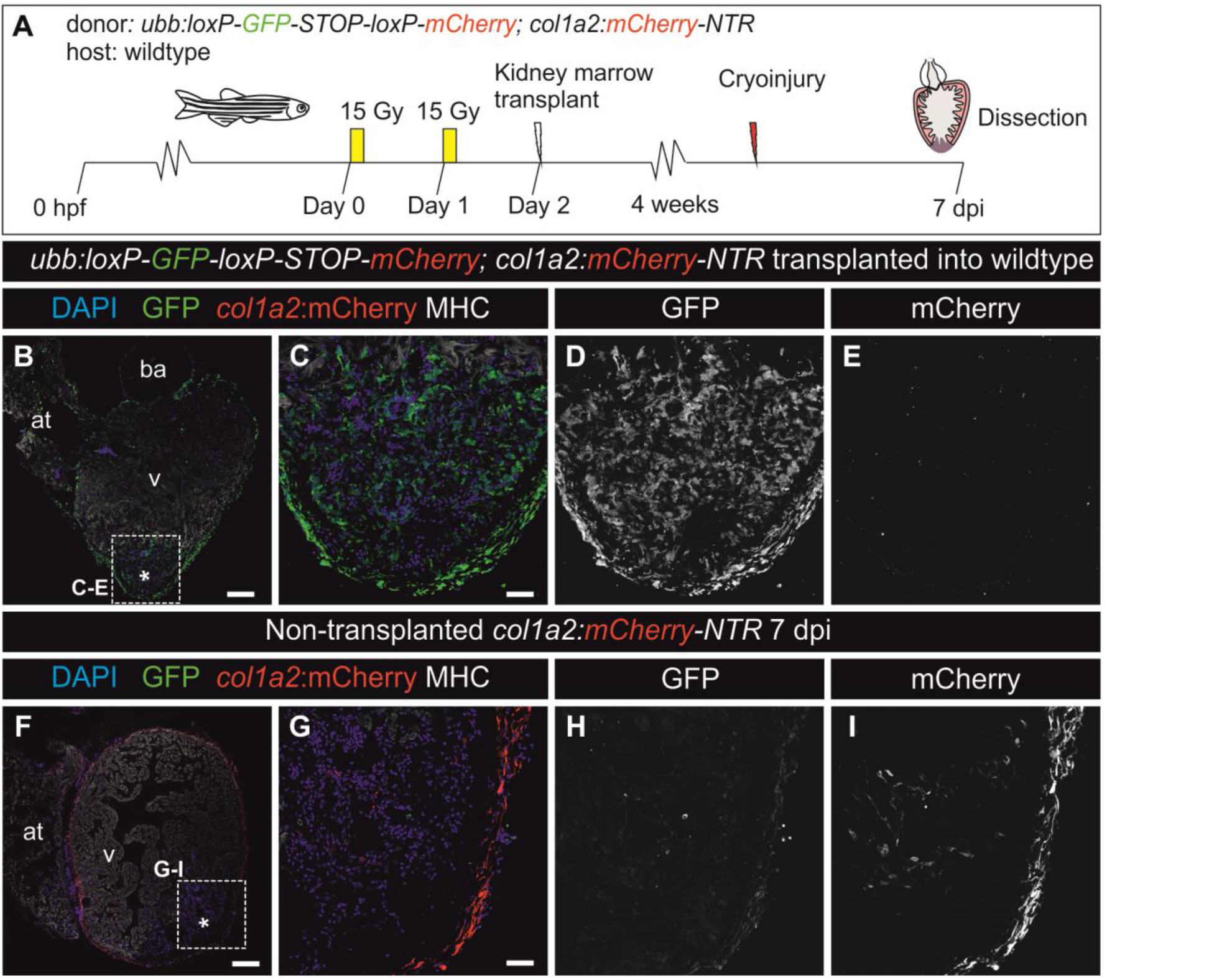
Kidney marrow derived cells do not contribute to collagen deposition during fibrotic response to cryoinjury. (*A*) Kidney marrow derived cells (KMDC) from a transgenic line ubiquitously expressing GFP and mCherry under the control of *col1a2* regulatory regions were transplanted into irradiated wildtype adult zebrafish. After kidney marrow reconstitution, host hearts were cryoinjured. Expression of mCherry and GFP was assessed at 7 days postinjury (dpi) by immunofluorescence on sections. Myosin Heavy Chain (MHC) marks myocardium, nuclei are counterstained with DAPI. (*B*) Whole heart section of a transplanted heart at 7 dpi. (*C-E*) Merged and single channels of boxed area in *B*. No mCherry signal is visible. (*F*) Control whole heart section from a *col1a2:mCherry* transgenic fish at 7 dpi. (*G–I*) Merged and single channels of boxed section shown in *F*; mCherry signal is detected in the epicardium and injury areas. Asterisks indicate injury area. Scale bars, 25 μm (*C* and *G*), 100 μm (*B* and *F*).

**Figure S4.**
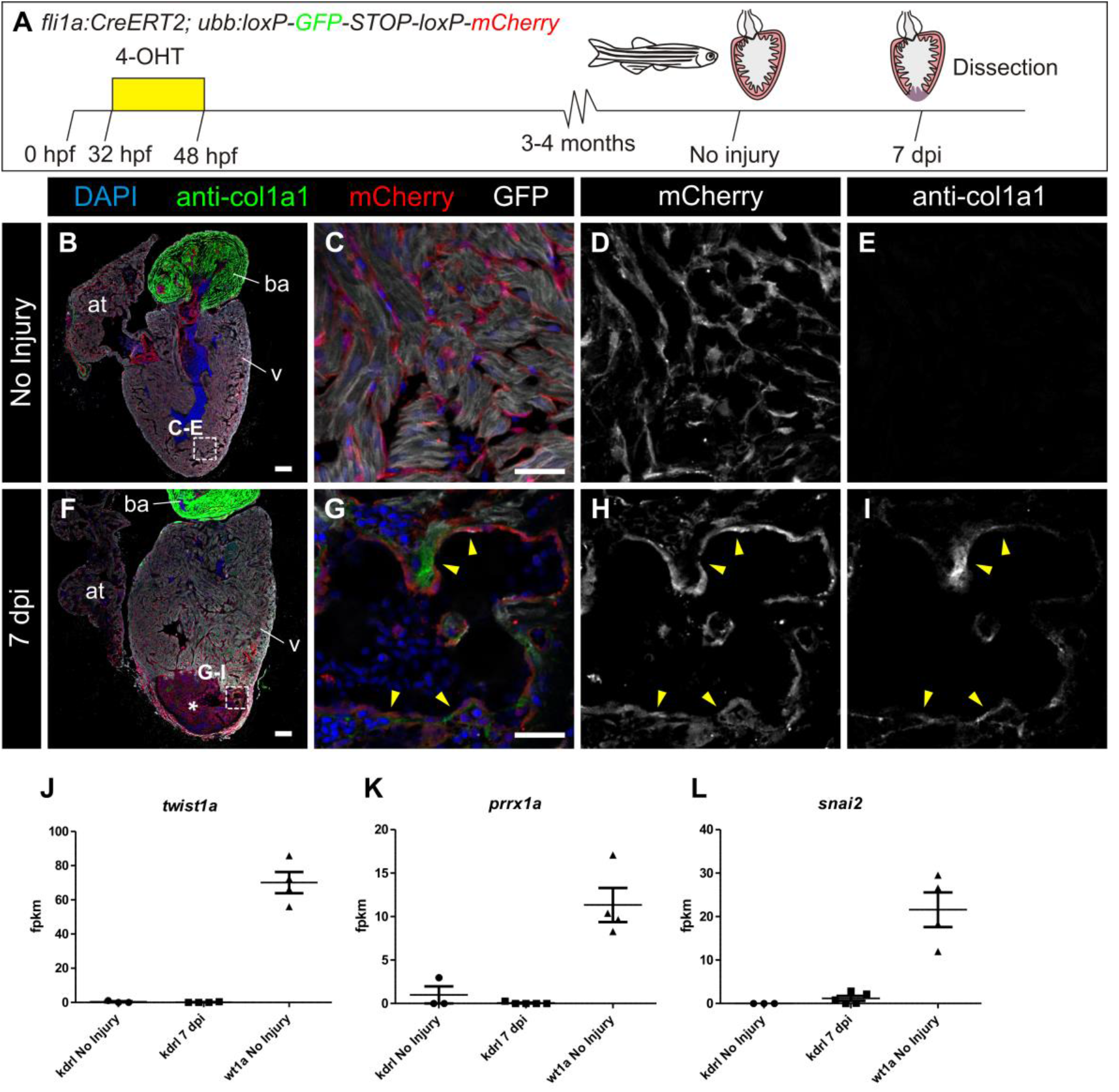
Endocardial cells at the injury area express collagen 1a1 but not EMT markers. (*A-I*) Lineage tracing of endocardial cells in the uninjured and injured heart. (*A*) Experimental scheme for tracing the contribution of endocardial cells to fibrosis using the *ubb:Switch* reporter line. (*B* and *F*) show immunofluorescence staining of whole-heart sagittal sections (*B*, uninjured; *F*, 7 dpi) with anti-mCherry (red), anti-col1a1 (green), and anti-Myosin Heavy Chain (MHC; grey); nuclei were DAPI counterstained (blue). Panels (*C–E*) and (*G–I*) are merged and single channels of a zoomed view of the endocardial border in the apex or close to the injury area (IA; asterisk). *fli1a*-derived cells are mCherry^+^ (red). Note that at 7 days postinjury (dpi), mCherry^+^ cells (red) are closely associated with col1a1 (green) deposits (arrowheads). (*J–L*) Epithelial to mesenchymal transition (EMT) markers are expressed by *wt1a*:GFP^+^ cells but not by *kdrl:mCherry*^+^ cells. Graphs show the fpkm values for the same genes and samples. EMT genes were more abundant in *wt1a*^+^ cells than in *kdrl*^+^ cells both before and after injury. at, atrium; ba, bulbus arteriosus; hpf, hours postfertilization; v, ventricle. Scale bars, 25 μm (*C, G*), 100 μm (*B, F*).

**Figure S5.**
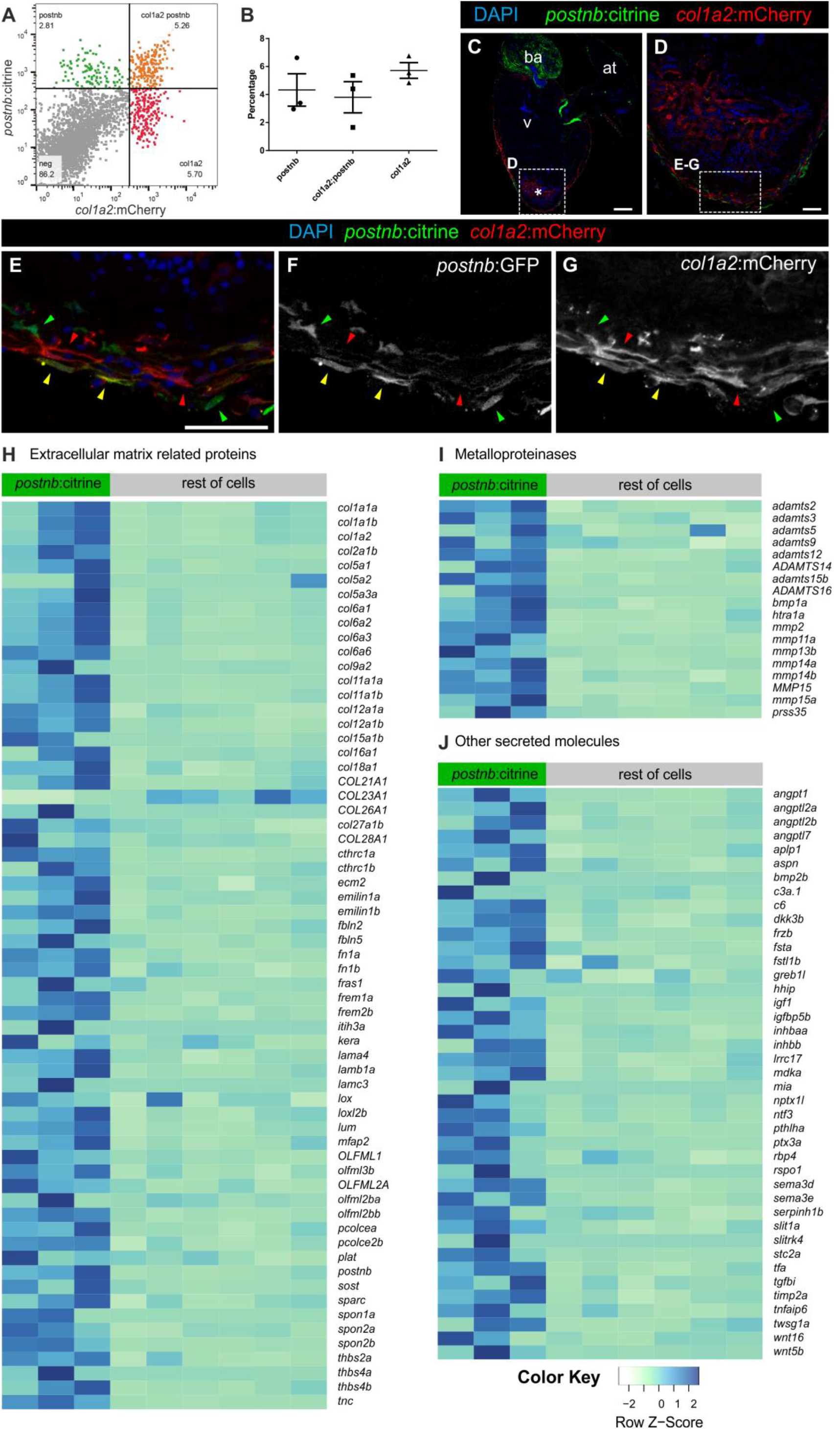
Colocalization of *postnb*:citrine^+^ and *col1a2*:mCherry^+^ cells. (*A* and *B*) FACS-sorted cells of 7 days postinjury (dpi) hearts from *co1a2:mCherry;postnb:citrine* double transgenic fish. Based on these markers, three populations could be detected: doublepositive, *postnb*^+^, and *col1a2*^+^. (*C–G*) Immunofluorescence with anti GFP (green) and anti-mCherry (red) on a heart section from an adult *postnb:citrine;col1a2:mCherry-NTR* fish. Note the presence of double-positive (yellow arrowheads), *col1a2*^+^ (red arrowheads), and *postnb*^+^ cells (green arrowheads). (*H–J*) Heatmap representing the expression levels of all the secreted proteins that were identified to be upregulated when comparing *postnb*:citrine^+^ cells at 7 dpi to the rest of cells at the injury (with the exception of the *kdrl*:mCherry population). fpkm values were used. The list of genes encoding secreted molecules was obtained by using Ingenuity software, and manually adding the genes encoding for collagens and metalloproteinases not identified by Ingenuity. For better visualization of changes in gene expression between both groups, values were scaled independently for each gene, that is, each row. at, atrium; ba, bulbus arteriosus; v, ventricle. Scale bars, 25 μm (*C*), 100 μm (D,G).

**Figure S6.**
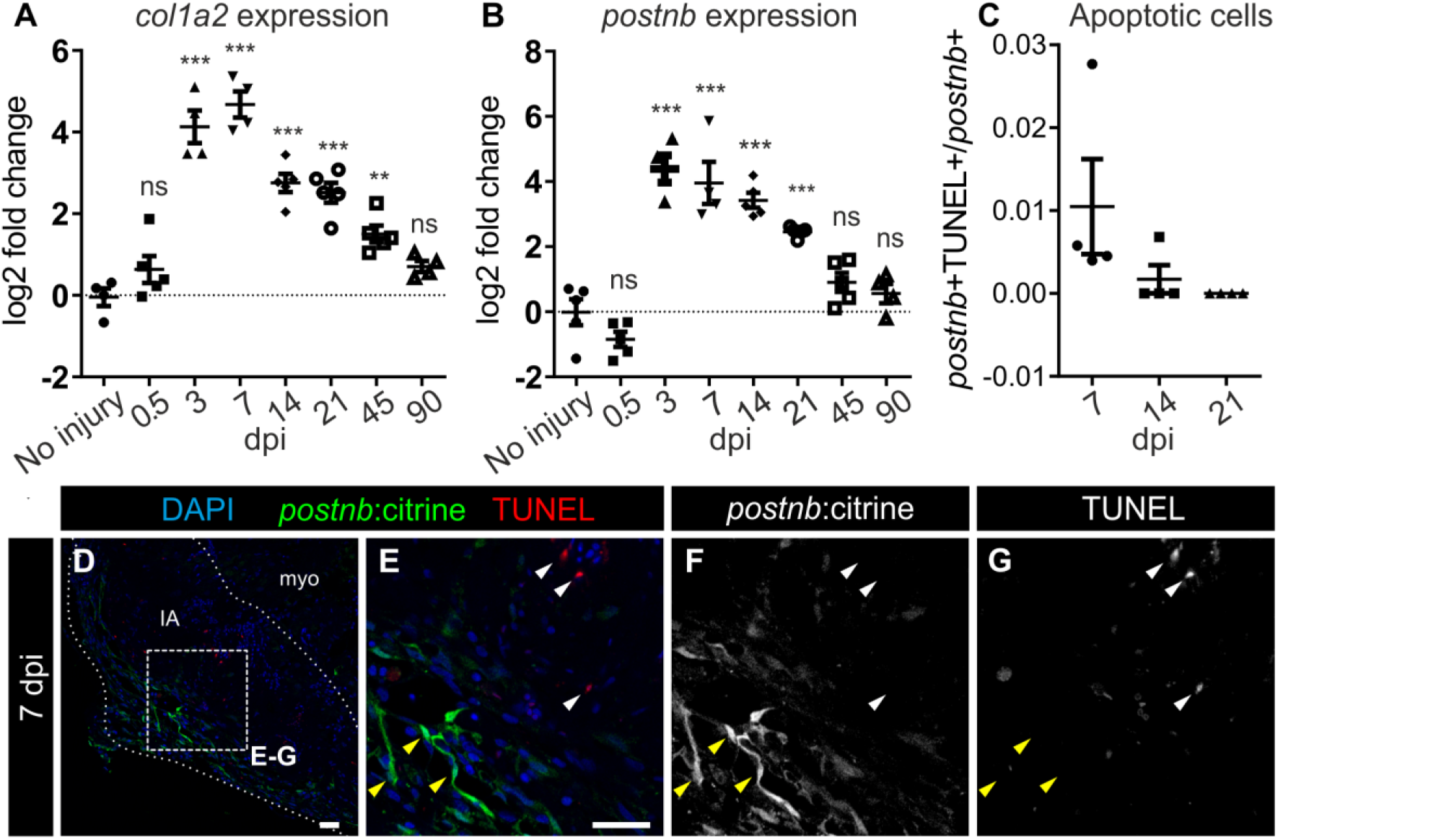
Apoptotic and senescent fibroblasts during heart regeneration. (*A* and *B*) qPCR of *col1a2* and *postnb* at different days postinjury. Symbols show data for individual samples, bars and whiskers show mean±s.d.; ^***^*P* < 0.001; ^**^*P* < 0.01 by one-way ANOVA followed by Tukey's multiple comparisons test. (C) *postnb*^+^ apoptotic (TUNEL^+^) cell quantification. (*D–G*) TUNEL staining on *postnb*:citrine heart sections. IA, injury area; myo, myocardium. Scale bars, 25 μm.

**Figure S7.**
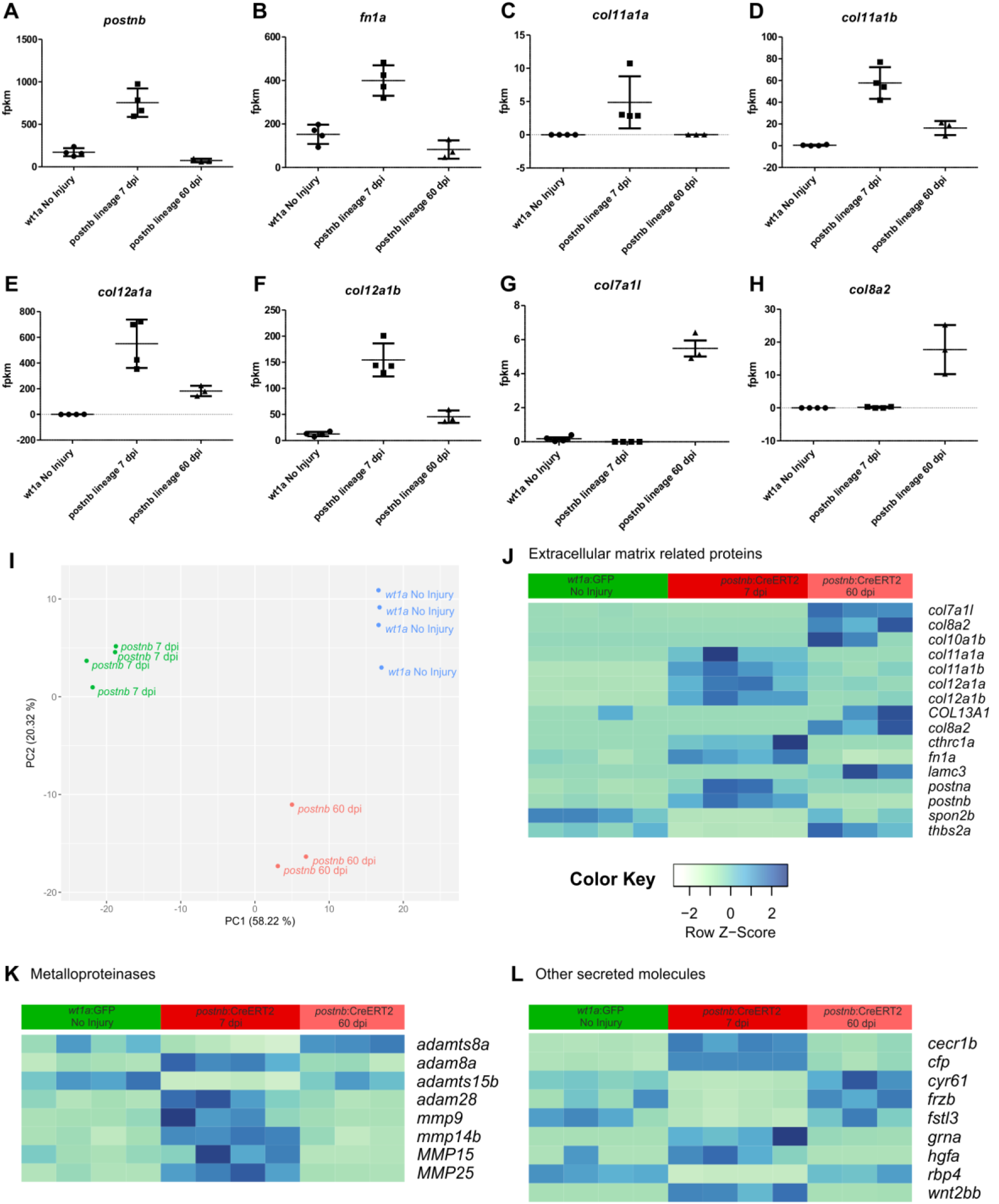
Comparison of the gene expression profile of resident intracardiac fibroblasts, activated fibroblasts, and inactivated fibroblasts. (*A–H*) Plots of fpkm values for the same genes and samples: extracellular matrix genes were more abundant in *postnb*-derived cells at 7 days postinjury (dpi). In some cases, the expression profile of *wt1a*:GFP and *postnb*-derived cells at 60 dpi did not coincide, suggesting that fibroblasts do not fully revert to a quiescent state during heart regeneration. The genes upregulated in *postnb*-derived cells at 60 dpi are expressed at low levels. (*I*) Principal Component Analysis of RNA-seq samples from uninjured *wt1a*:GFP^+^ cells and postnb-derived cells from hearts at 7 dpi and 60 dpi. (*J–L*) Heatmap representing the expression levels of all the secreted proteins that were identified to be differentially expressed when comparing *postnb*:CreER^T2^ lineage traced cells at 7 dpi to those cells at 60 dpi (with the exception of the *kdrl*:mCherry population). Expression levels in the *wt1a*:GFP population are also shown for comparison. fpkm values were used. The list of genes encoding for secreted molecules was obtained by using Ingenuity software, and manually adding the genes encoding for collagens and metalloproteinases not identified by Ingenuity. For better visualization of changes in gene expression between both groups, values were scaled independently for each gene, that is, each row.

**Figure S8.**
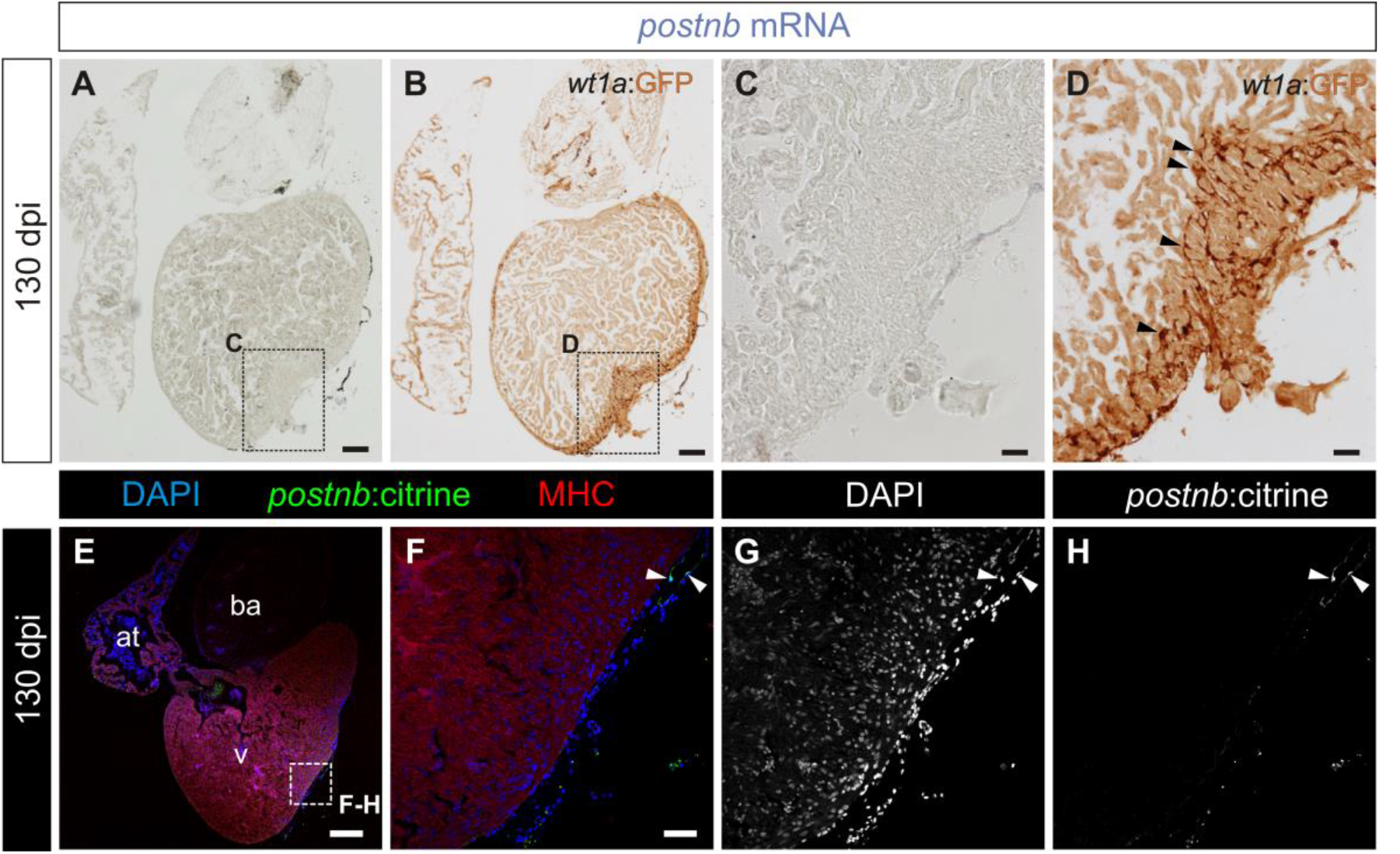
Expression pattern of *wt1a*:GFP and *postnb*:citrine in fully regenerated hearts (*A–D*) *postnb* mRNA *in situ* hybridization (purple) followed by anti-GFP immunohistochemistry (brown) on sections of *wt1a:GFP* ventricles at 130 days postinjury (dpi) (n = 3/3). Arrowheads mark *wt1a*:GFP^+^ cells. Sections were stained simultaneously with those shown in Figure 1 (*C–F*). (*E–H*) Immunofluorescence with anti-GFP (green) and anti-Myosin Heavy Chain (MHC) (red) on a *postnb:citrine* heart section at 130 dpi (n = 4/4). Nuclei are counterstaind with DAPI. Arrowheads mark GFP^+^ cells. at, atrium; ba, bulbus arteriosus; v, ventricle. Scale bars, 100 μm (*A* and *B*), 25 μm (*C* and *D*).

**Figure S9.**
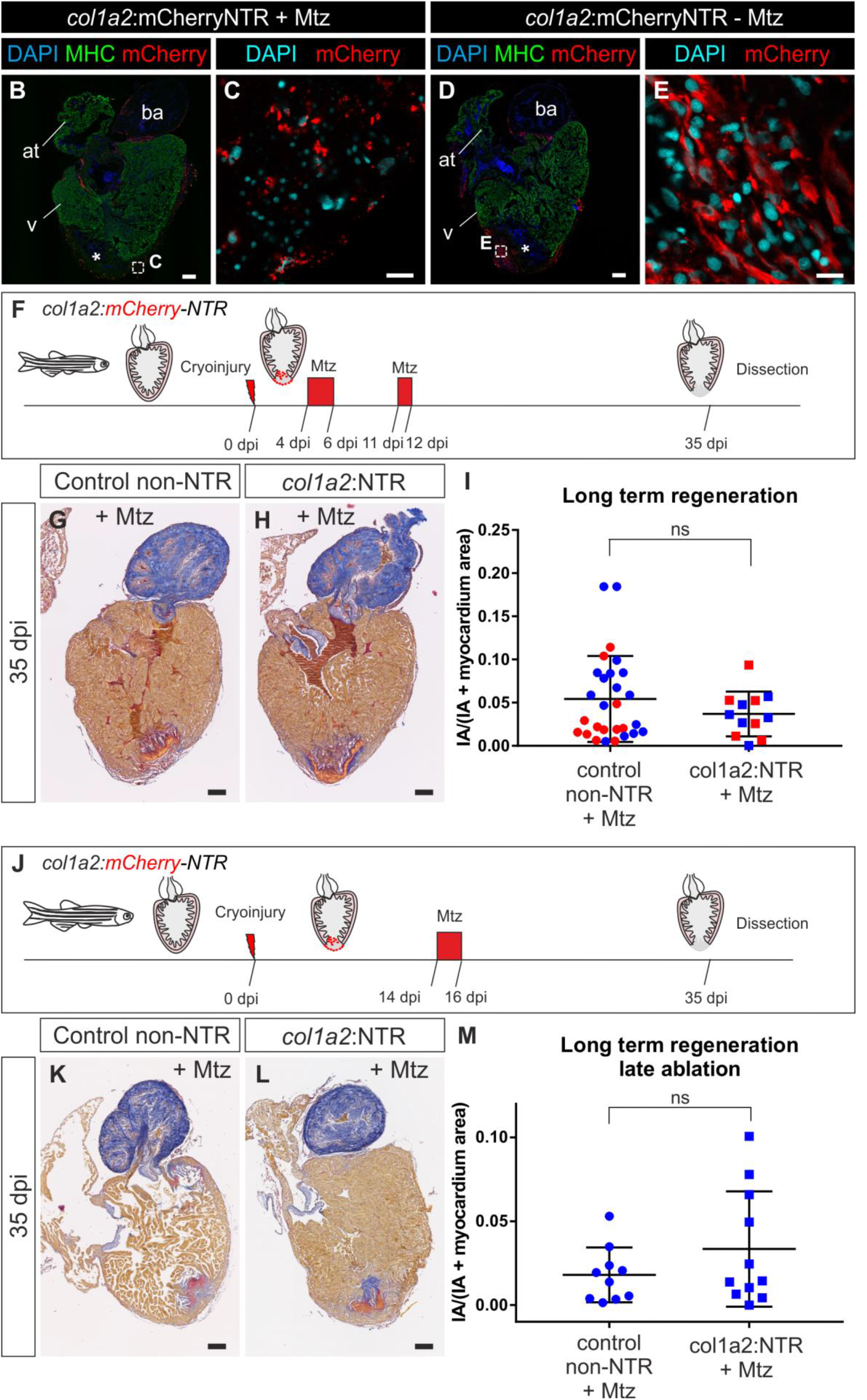
Heart regeneration upon genetic ablation of *col1a2*:expressing cells. (*A*) *col1a2*:mCherry-*NTR* (in short *col1a2:NTR*) adult animals were cryoinjured and treated with Metronidazole (Mtz) from 4 to 6 days postinjury (dpi). BrdU injection was performed one day prior to fixation to assess cardiomyocyte proliferation. (*B–E*) Immunofluorescence on heart sections of *col1a2:mCherry-NTR* treated with Mtz (*B* and *C*) or untreated controls (*D* and *E*). mCherry, red; myosin heavy chain (MHC), green; nuclei (DAPI), blue for (*B*) and (*D*), and cyan for (*C*) and (*E*). In Mtz-treated fish, *col1a2*:mCherry-NTR labels cells with fragmented nuclei and the homogeneous mCherry expression observed in the wildtype heart is lost. (*F*) Experimental scheme to study regeneration upon genetic ablation of col1a2 expressing cells. *col1a2:NTR* transgenic zebrafish were cryoinjured and treated with 10 mM Metronidazole (Mtz) between 4 to 6 and 11 to 12 days postinjury (dpi). Hearts were dissected at 35 dpi, sectioned and stained with AFOG to determine degree of regeneration. (*G* and *H*) AFOG-stained sagittal section through ventricles of a Mtz-treated *col1a2:loxP-tag SFP-loxP-mCherry-NTR* heart (control) and a Mtz-treated *col1a2:NTR* heart. (*I*) Quantification of the injury area versus total ventricular area from 28 control hearts and 12 *col1a2:NTR* hearts. Blue and red colors indicate results from two independent experiments. No significant difference was observed between control and *col1a2*:NTR groups by unpaired Student's t-test (*P* = 0.75). (*J*) Experimental scheme. *col1a2:NTR* transgenic zebrafish were cryoinjured and treated with 10 mM Mtz between 14 and 16 dpi. Hearts were dissected at 35 dpi, sectioned and stained with AFOG to determine degree of regeneration. (*K* and *L*) AFOG-stained sagittal section through ventricles of a Mtz-treated *col1a2:loxP-tag SFP-loxP-mCherry-NTR* heart (control) and a Mtz-treated *col1a2:NTR* heart. (*M*) Quantification of the injury area versus total ventricular area from 10 control hearts and 11 *col1a2:NTR* hearts. No significant difference was observed between control and *col1a2*:NTR groups by unpaired Student's t-test (*P* = 0.2113). IA, injury area. Scale bars, 100 μm.

**Figure S10.**
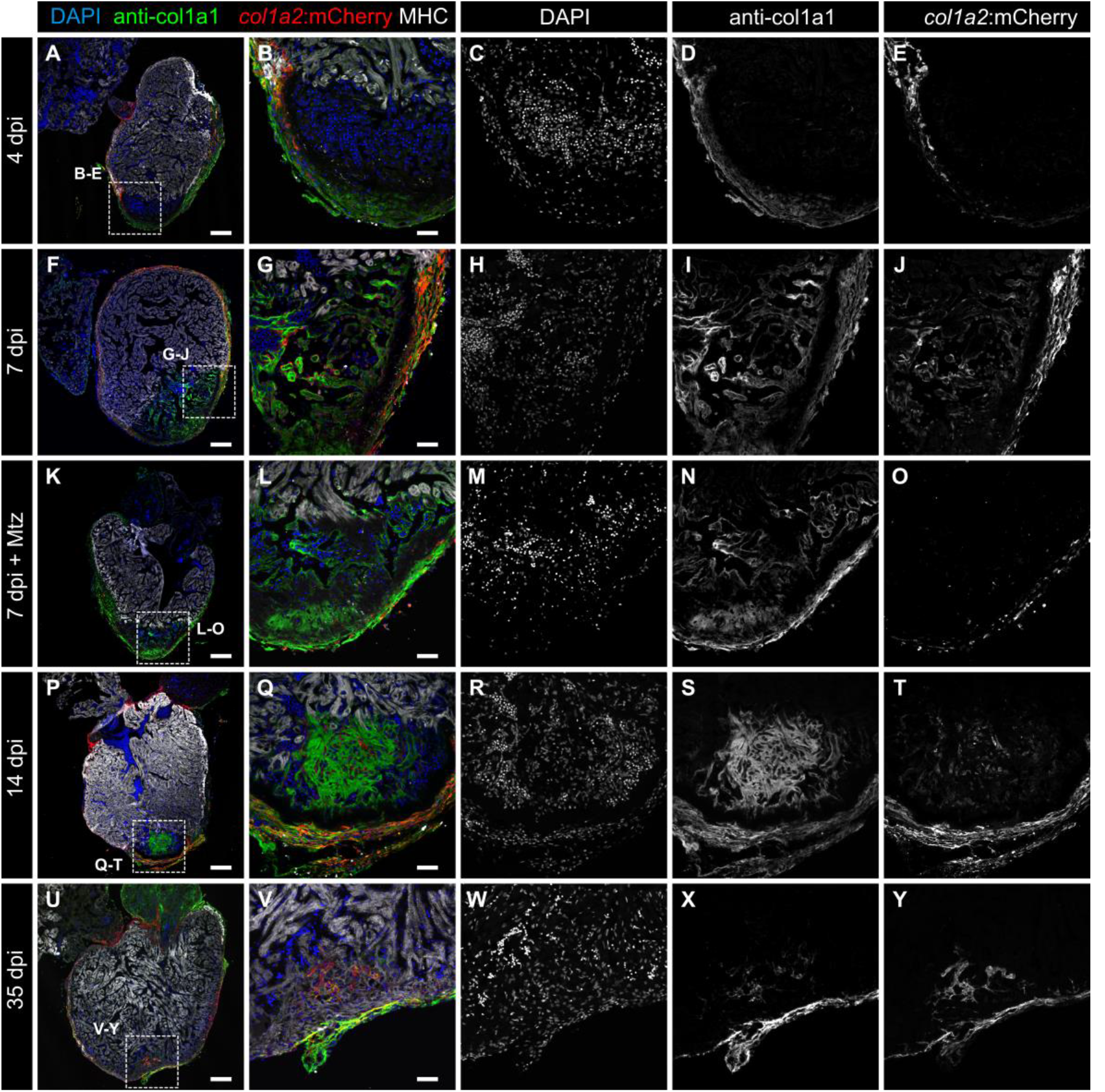
Colocalization of *col1a2*:mCherry and anti-col1a1 at different stages post injury and ablation. (*A–Y*) Sagittal sections through *col1a2:mCherry* hearts at 4 days post injury (dpi) (*A–E*; n = 4/4), 7 dpi (*F–J*; n = 4/4), 7 dpi treated with Metronidazole (Mtz) (K–O; n = 4/4), 14 dpi (P–T; n = 4/4) and 35 dpi (U–Y; n = 3/3). Sections were immunostained with anti-mCherry (red), anti-col1a1 (green) and anti-Myosin Heavy Chain (MHC, white). Nuclei are counterstained with DAPI (blue). Scale bars, 25 μm (*B, G, L, Q, V*), 100 μm (*A, F, K, P, U*).

## SUPPLEMENTARY MATERIALS AND METHODS

### Transgenic lines

The construct to generate Tg(*postnb*:citrine)^cn6^ was obtained by recombining the iTol2Amp cassette (1) followed by the citrine-Kan cassette into the BAC CH211-38D6. The construct for Tg(*periostin:CreER^T2^*)^cn7^ was generated recombining the *iTol2Amp-Cryst:GFP* cassette followed by *CreERT2-frt-Kan-frt* into the BAC CH73-370H18. The plasmid to generate Tg(*col1a2:loxP-tag SFP-loxP-mCherry-NTR*)^cn8^ was obtained by recombining the *iTol2-CrystCFP* (2) and *loxP-tag SFP-loxP-mCherry-NTR* cassettes into the BAC CH211-122K13. Once the transgenic line was established, *Cre recombinase* mRNA was injected at 50 ng μl^-1^ to recombine the *loxP* sites to generate the *Tg(col1a2:loxP-mCherry-NTR*)^cn11^.

The plasmid to generate the Tg(*fli1a:CreER^T2^*)^cn9^ line was obtained using Gateway (3) to recombine the plasmids p5E-fli1ep (4), pME-CreER^T2^ (5), p5E-pA (3) into the pDestTol2pA2 (4) backbone.

Plasmid templates for recombineering and the plasmid to generate the Tg(*wt1a*:CreER^T2^)^cn10^ line were cloned using Gibson Assembly (NEB) using the promoter reported previously (6). Recombineering was performed combining pRedET (GeneBridges, Germany) system and EL250 bacteria (7). DNA was injected at 25 ng μL^-1^ into one-cell stage zebrafish embryos along with 1 nl of 50 ng μL^-1^ synthetic *Tol2* mRNA in Danieau buffer.

Other lines used were Tg(*kdrl:mcherry*) (kindly provided by E. Ober), Tg(-3.5*ubb:loxP--loxP-eGFP*)^cz1702Tg^ (5), Tg(*ubb:Switch*)^cz1701^ (5), Tg(*ubb:mCherry*)^cz1705Tg^ (5), Tg(*fli1a:GFP*)^y1^ (8) Tg1 (*-6.8wt1a:EGFP*)^li7Tg^ (6), and Tg(*tcf21:CreER^T2^*)^pd42Tg^ (9).

In adults, 10 μM 4-OHT was administered at the indicated times dissolved in 100 mL of fish water per fish. Treatments were performed overnight for 12 hours. Prior to administration, the 10 mM stock (dissolved in ethanol) was heated 10 minutes at 65 °C (10). In embryos, it was administered at 5 μM. Cryoinjury was performed as previously described (11). Genetic ablation was performed using the Nitroreductase system previously described (12). Metronidazole (Sigma) was added at 10 mM to fish water at the indicated times.

### Whole kidney marrow transplantation

Transplantations were performed as described (13).

### Histology and sectioning

Samples for Fig. 1 C–F, Fig. 5, Fig S 4, Fig. S 6, Fig. S 8 and Fig S 9 G-M were fixed in 4% paraformaldehyde (PFA) in phosphate buffered saline (PBS) overnight at 4°C, washed in PBS + 0.1% Tween 20 (Sigma), dehydrated through an ethanol series and embedded in paraffin. They were sectioned at 7 μm with a microtome (Leica), mounted on Superfrost slides (Fisher Scientific) and dried overnight at 37°C. Sections were deparaffinized in xylol, rehydrated and washed in distilled water. Connective tissue was stained using Acid Fuchsin Orange G (AFOG). For analysis of the amount of IA/IA + myocardium (Fig. S8), the total ventricular tissue area and IA on all sections on a slide of each heart (collected on 5 slides) was measured.

Other sections were processed by fixing in 4% PFA and washing in PBS + 0.1 % Tween 20. They were then incubated in 15% saccharose overnight 4°C, embedded in 30% gelatin 15% saccharose and snap frozen at -80 °C in isopentane. They were cut at 8 μm on a cryostat (Leica).

For immunofluorescence, whole mount hearts were fixed in 4% PFA overnight, washed in PBS + 0.1% Tween and permeabilized with PBS + 0.5% Triton-X100 (Sigma) for 20 min. Several washing steps were followed by at least 2 h of blocking with 5% goat serum, 5% BSA, 20 mM MgCl_2_ in PBS, and then slides were incubated with the antibodies overnight.

Paraffin sections were deparaffinized, rehydrated and washed in distilled water. Epitopes were retrieved by heating in 10 mM citrate buffer (pH 6.0) for 15 minutes in a microwave oven at full power. Gelatin sections were incubated for 30 min in PBS + 0.1% Tween 20 at 37°C to dissolve the gelatin. Non-specific binding sites were saturated by incubation for at least 1 h in blocking solution (as above). Endogenous biotin was blocked with the avidin-biotin blocking kit (Vector, Burlingame, CA, USA).

In situ hybridization was performed using *col1a2 and postnb* riboprobes (14).

Primary antibodies used were anti-myosin heavy chain (MF20, DSHB, 1:20 and F59, DSHB, 1:20), anti-GFP (AVES, GFP-1010, 1:500; Clontech, 632592, Mountain View, CA, USA, 1:100), anti-col1a1 (SP1.D8, DSHB, 1:20), anti-RFP (ab34771, Abcam, 1:200), anti-Mef-2 (Santa Cruz Biotechnology, C21, sc-313, 1:200), anti-BrdU (BD Biosciences, B44, 1:100) Biotin- or Alexa (488, 568, 633) -conjugated secondary antibodies and streptavidin-Cy3 (Jackson Immuno Research Laboratories) were used at 1:300. Nuclei were stained with DAPI and slides were mounted in Fluorsave (Calbiochem).

### Quantitative real-time (qRT) PCR

RNA from cardiac ventricles was extracted using 0.5 mL Trizol reagent (Ambion, Life Technologies). One ventricle was used per biological replicate. RNA was transcribed to cDNA using the High-Capacity cDNA Reverse Transcription Kit (Applied Biosystems, Life Technologies). qRT-PCR was performed using Power SYBR Green PCR Master Mix (Applied Biosystems). *col1a2* and *postnb* expression was normalized with the geometric mean of the expression level of two constitutive genes, *EF1-alpha* and *rps11*.

### Heart dissociation, sorter and RNA-Seq library production

For Tg(*kdrl:mCherry; postnb:citrine*) fish, the apex of the heart was dissected, and dissociated for 45 min in trypsin. Whole ventricles (for Tg(*wt1a:GFP*)) or apex-for Tg(*postnb:CreERT2;ubi:Switch*)- were dissected and dissociated according to previous protocols (15) with minor modifications: the enzyme concentration was doubled and time of digestion increased to 2 h. Then, one volume of PBS + 10% Fetal Bovine Serum (FBS) was added to the sample, which was centrifuged for 8 min at 250 × g, and re-suspended in PBS + 1% FBS.

### Cell sorting and RNA-seq library production

Cells were sorted on a SONY Synergy sy3200 cell sorter (Sony Biotechnology) and RNA was extracted using the Arcturus Pico Pure kit (Thermofisher). Pools of 3–5 apex/ventricles were used.

For *kdrl:mCherry*^+^ cells, three pools of 3–5 hearts were used for the uninjured condition and 5 pools of 3–5 hearts were used for the 7 dpi condition. Three pools of 3–5 ventricular apices were used for postnb:citrine cells per condition, and 6 pools of 3–5 ventricular apices were used for the remainder of cells. For *postnb*-derived cells, four pools of 3–5 hearts were used for 7 dpi, and 3 pools were used for 60 dpi.

Total RNA (0.25–1 ng) was used to generate barcoded RNA-seq libraries using the Ovation Single Cell RNA-Seq System (NuGEN) with two rounds of library amplification. The size of the libraries was calculated using the Agilent 2100 Bioanalyzer. Library concentration was determined using the Qubit^®^ fluorometer (ThermoFisher Scientific). Libraries were sequenced on the HiSeq2500 system (Illumina) to generate 60 bases single reads. FastQ files for each sample were obtained using CASAVA v1.8 software (Illumina). Four biological replicates consisting of 5 pooled hearts were used per sample.

### RNA-Seq analysis

Sequencing adaptor contaminations were removed from reads using cutadapt 1.9.1 software (16) and the resulting reads were mapped and quantified on the transcriptome (Ensembl gene-build 10, release 82) using RSEM v1.2.25 (17). Only genes with at least 1 count per million in at least 2–3 samples were considered for statistical analysis. Data were then normalized and differential expression tested using the bioconductor package EdgeR (18). We considered as differentially expressed those genes with a Benjamini-Hochberg adjusted *P*-value ≤0.05 and LFC ≥ 1. For the Tg(*wt1a:GFP*) samples, paired analysis was used. Heatmaps were made using gplot library and heatmap.2 function.

